# A Computational Pipeline for Glioblastoma Vaccine Development: Integrating Novel Omics-Driven OIP5 Target Discovery to Create a Deep Learning-Based Immunogenicity Framework for Personalized Immunotherapy

**DOI:** 10.64898/2025.12.23.696316

**Authors:** Rahul Mahapatra, Anish Sundar, Aarsh Shekhar

## Abstract

This work introduces a modular, open-source computational pipeline for glioblastoma (GBM) vaccine design that integrates omics-based OIP5 target discovery with a deep learning framework for epitope immunogenicity prediction. Building upon conventional affinity-based predictors such as NetMHCpan, our Tumor Epitope Immunogenicity Pipeline (TEIP) incorporates biological, structural, and transcriptomic context to predict tumor-specific T-cell responses. Using curated immunogenic and non-immunogenic peptides from IEDB and GBM datasets, TEIP employs dual bidirectional LSTM encoders to represent peptide and HLA sequences, concatenated with auxiliary molecular features including proteasomal cleavage, TAP transport likelihood, gene expression, and mutation frequency. These features are fused into a probabilistic model that outputs peptide-specific immunogenicity scores and confidence estimates. Benchmark results across multiple HLA alleles demonstrate that TEIP outperforms NetMHCpan and MARIA, achieving an ROC-AUC of 0.89 and PR-AUC of 0.35. Feature importance analysis confirms that tumor-specific expression and peptide-MHC binding dominate predictive accuracy, validating the biological realism of the model. By combining mechanistic features with data-driven representation learning, TEIP enables fine-grained prioritization of neoantigens with translational potential. In proof-of-concept GBM studies, TEIP recovered known cancer-testis antigens such as OIP5, demonstrating its ability to identify clinically relevant vaccine targets. The pipeline is implemented using entirely open-access data and software to promote reproducibility and scalability. Collectively, TEIP provides a unified framework that connects multi-omics tumor characterization with deep learning–based antigen modeling, establishing a foundation for precision immunotherapy development in glioblastoma and beyond.

## I. Introduction

Glial cell are non-neuronal supportive cells in the central nervous system that provide protection and nourishment for neurons [1]. Gliomas, tumors of glial tissue, can cause a wide variety of symptoms depending on their location in the central nervous system. Out of the many kinds of gliomas, not all are malignant, with some kinds growing slowly and not considered cancerous, and some growing quickly and invading brain tissue [2]. The most common type of gliomas are astrocytomas, which evolve from star shaped glial cells known as astrocytes and account for about half of malignant brain tumors.

Astrocytomas are classified by WHO from grades 1-4 in terms of tumor growth. Grade 1 and 2 astrocytomas are both considered low grade, but grade 2 astrocytomas have higher coverage than grade 1. Grade 3 and 4 astrocytomas are considered high grade, and tend to spread rapidly. Until quite recently, the most aggressive astrocytoma, glioblastoma multiforme, was considered a grade 4, but now, according to the 2021 WHO Classification of Central Nervous System Tumors, glioblastoma (GBM) is recognized as a distinct molecular entity characterized by IDH wildtype status, TERT promoter mutations, EGFR amplification, and/or chromosome 7 gain with chromosome 10 loss (7+/10–) [3][4].

Around 46.1% of all malignant primary brain tumors are glioblastomas, underscoring its prevalence and clinical importance [5]. Despite decades of research, the median survival stays stagnant at only 12–18 months even with full treatment: surgical resection followed by concurrent radiotherapy and temozolomide chemotherapy [6]. Recurrence has been documented repeatedly, and no current therapy offers a cure [7].

One of the major obstacles to effectively countering the effects of GBM is its immunologically “cold” tumor environment, which is dominated by immunosuppressive cells such as tumor associated macrophages and microglia [7][8]. Moreover, T-cell infiltration is rather sparse in the target area. The blood-brain barrier (BBB) inhibits drug delivery to a certain extent, and the tumor’s high heterogeneity allows any resistant subclones to survive after therapy [9]. Overall, these factors explain why conventional methods of treatment like targeted kinase inhibitors and immune checkpoint blocking therapies have failed to yield beneficial effects against GBM.

Because of these barriers, novel immunotherapeutic strategies are a pressing issue concerning GBM’s ability to evade immune responses. A prospect for treatment involves targeting cancer-testis antigens (CTAs), which are a class of proteins that are typically only expressed in germline tissues, but are aberrantly activated in various cancers [10]. CTAs are especially promising because they are absent in normal somatic cell tissues, meaning that off-target toxicity characteristic of cancer therapy is minimized [11][12]. In addition, their restricted expression in the immune-privileged sites of the testis means that the immune system is bound to “think” of the aberrantly expressed CTA as foreign. Because this new exposure is considered a threat by the immune system, it can create a highly pointed attack when CTAs are presented on cancer cells[12].

Among CTAs, we have found Opa Interacting Protein 5 (OIP5) to be an especially compelling candidate for novel advancements in GBM therapy. OIP5 plays an essential role in centromere assembly and cell division, and its expression is highly elevated in glioblastoma tissues compared to normal brain tissue, where its presence is mainly undetectable [13][14]. Studies have shown that knockdown in glioma cell lines lead to cell cycle arrest and apoptosis, thus confirming its potential critical role in tumor proliferation [13]. OIP5’s immunogenic potential means that it could serve as a target for T-cell based therapies such as peptide vaccines or T-cell receptor engineered therapies (TCR-T), which are designed to activate certain cytotoxic T lymphocytes against GBM cells that express epitopes derived from OIP5 [15][16].

## II. Research Rationale

Existing literature has predominantly examined OIP5’s role in chromosomal segregation and tumor cell cycle control. However, its immunogenic landscape or feasibility as a candidate for vaccine construction is not deeply studied. This gap in research presents an opportunity to evaluate OIP5 through a computational immunology framework, in which multi-omic tumor data with peptide-HLA prediction tools will be utilized to gauge its value as a tumor-specific antigen for GBM.

The rationale behind this study is to leverage in silico analysis to validate whether OIP5 fulfills the criteria for a viable T-cell target: (1) strong overexpression in GBM relative to normal brain tissue, (2) limited or absent expression in vital tissues to minimize toxicity, (3) essential function in tumor proliferation to reduce escape potential, and (4) presence of short peptide fragments that can stably bind to common HLA class I molecules and be recognized by cytotoxic T cells. If OIP5 fulfills these criteria, it could act as the foundation for a vaccine or adoptive cell therapy with the potential to create durable, tumor-specific immune responses.

This project also addresses the major challenge in GBM immunotherapy: heterogeneity and patient variability. Each patient’s tumor may differ in antigen expression and HLA type. Therefore, our study proposes a multi-epitope approach, in which several high-affinity OIP5 derived peptides restricted by HLA alleles (e.g., A02:01, A24:02, B^*^07:02) will be identified. This redundancy ensures that even if certain epitopes are not processed or presented efficiently, others remain immunogenic across a broad population. Moreover, combining the epitope prediction with population coverage modeling will estimate the proportion of patients likely to benefit from this vaccine design.

Our entirely computational workflow creates a reproducible and cost-effective route to early-stage immunotherapy design. This research uses open-access databases like TCGA, GTEx, and DepMap for expression and dependency validation. Moreover, tools such as IEDB, NetMHCpan, and AlphaFold will be incorporated for epitope modeling. Beyond the topic of glioblastoma, this approach can be generalized to other cancers that aberrantly express CTAs, which can accelerate the identification of novel immunogenic targets while lowering laboratory costs and ethical boundaries.

Ultimately, this study aims to bridge the gap between bioinformatics and translational immunology. It provides a first step to an OIP5 targeted vaccine for GBM and offers a proofof-concept for how computational biology can systematically guide immunotherapy development.

## III. Methodology

A proposed solution is to design a multi-epitope peptide vaccine that can train cytotoxic T cells to recognize and eliminate glioblastoma cells that aberrantly express OIP5. This methodology utilizes the ability of the immune system to infiltrate the central nervous system (CNS), providing an advantage over conventionally used drugs that are obstructed by the blood-brain barrier. The vaccine will consist of several short OIP5 peptides that are predicted to bind strongly to several common HLA class I molecules. By including epitopes from different regions of the OIP5 protein, this experimental design accounts for the intratumoral heterogeneity and broad patient coverage required for glioblastoma.

All the stages of vaccine development will be done entirely in silico using open-access bioinformatics tools and datasets. The computational workflow begins validation of OIP5 as a tumor-specific, essential gene by using TCGA, GTEx, GEPIA, and DepMap datasets. Next, epitope prediction will be completed using predictive softwares like IEDB and NetMHCpan. In the process, these tools will identify 8-11 amino acid fragments from OIP5 that are predicted to bind to common HLA class I alleles that have high affinity. These prospective epitopes will be evaluated further for proteasomal processing likelihood, TAP transport efficiency, and then absence of homology to self-peptides to prevent reactivity that affects offtarget tissues. Then, modeling tools like AlphaFold, Rosetta, and molecular dynamics simulations will be used to confirm stable peptides to MHC binding and validate immunogenic potential.

In order to address patient-specific variability, population coverage analysis will be done using IEDB’s Population Coverage tool to estimate what percentage of patients would likely present at least one of the chosen peptides. The multiepitope formatting of the vaccine will provide redundancy against antigen loss or mutation. This ensures that if one peptide is not expressed or processed efficiently, others still remain immunogenic. In future repetitions, the same pipeline used here could be customized for individual patients by utilizing each tumor’s HLA genotype and RNA expression profile. Thus, this would enable a fully personalized vaccine. The final vaccine construct will integrate the selected OIP5 peptides with a confirmed adjuvant to enhance T-cell training and generate durable antitumor immunity.

## IV. Target Validation of OIP5 as a CTA in Glioblastoma

### A. OIP5 Expression in Glioblastoma vs. Normal Brain

In order to confirm that OIP5 is transcriptionally upregulated in glioblastoma, differential expression analysis was completed using GEPIA2 [17]. This tool integrates RNA seqdata from TCGA and GTEx cohorts. Results revealed that OIP5 mRNA expression was significantly higher in GBM tissues (n = 163) compared to normal brain tissues (n = 207) as shown in (Fig. 1). The median expression level in tumor samples was several times greater than in normal samples, indicating that OIP5 activation is a recurring feature of GBM. This observation supports the hypothesis that OIP5 contributes to tumor proliferation and therefore establishes its relevance as a prospective immunotherapy target.

**Fig. 1.**
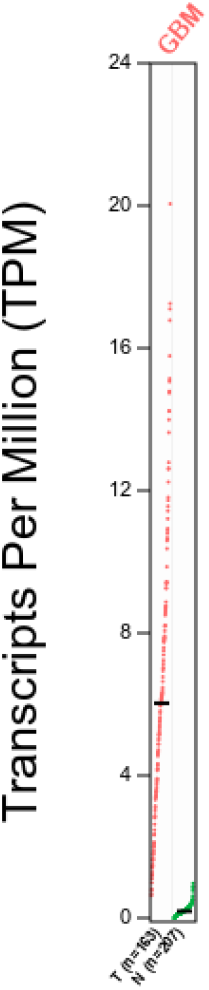
Comparison of OIP5 expression between normal brain and GBM tissue using GEPIA2. Tumor samples (red) exhibit markedly elevated expression relative to normal samples (green).

### B. OIP5 Displays Cancer-Testis Antigen Expression Patterns (GTEx Analysis)

Upon conducting analysis of the GTEx dataset, it was found that OIP5 expression in normal tissues is almost exclusively restricted to the testis [18]. There is minimal expression in other somatic tissues, including the brain (Figure 2). This restricted distribution pattern is characteristic of cancer-testis antigens (CTAs). These genes are normally active only in immune-privileged germline tissues. The low baseline expression of OIP5 in normal brain tissue along with its reactivation in glioblastoma clearly demonstrates that OIP5 meets the key requirement for CTA-based immunotherapy. It has tumor specific expression with minimal off-target risk.

**Fig. 2.**
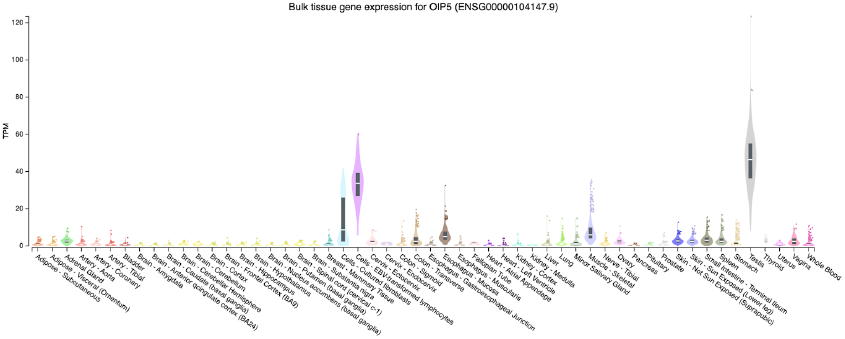
OIP5 tissue-wide expression profile (GTEx). Bar graph showing OIP5 mRNA expression across 30+ normal tissues, with a clear testis-dominant pattern and negligible expression in brain tissue.

### C. OIP5 Copy Number and Mutation Landscape in GBM (cBioPortal Analysis)

Data from cBioPortal (TCGA-GBM cohort) were used to assess genomic alterations that affect OIP5 [19]. As shown in Figure 3, OIP5 mRNA expression is positively correlated with copy-number gain, while tumors with shallow or deep deletions display reduced expression. Most GBM samples remain diploid or copy-number gain and deep deletions are rare. These results suggest that OIP5 is retained and possibly essential for tumor proliferation. Mutation frequency within the OIP5 locus was minimal (¡1%), further validating that OIP5 is not commonly inactivated in GBM. The genomic stability and expression independence hint that tumor cells may rely on OIP5 function, thus making it a stable immunotherapeutic target.

**Fig. 3.**
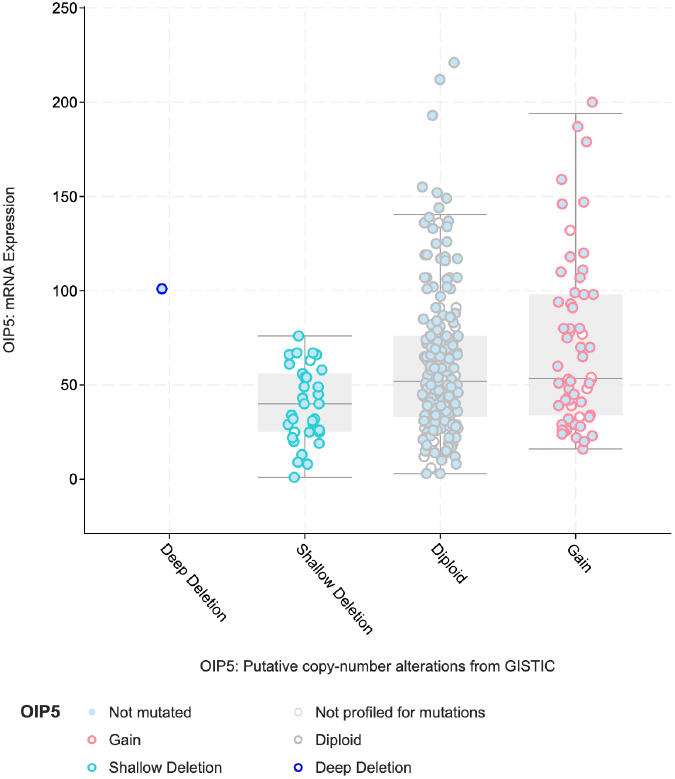
OIP5 mRNA expression level and OIP5 copy number categories for 281 samples in cBioPortal. In the dataset, OIP5 shows high mRNA expression in samples with copy number gain and a little amount of deletions. Most of the samples appear to be not mutated as well.

### D. OIP5 Dependency analysis using DepMap

Using the CRISPR (DepMap Public 25Q3+Score, Chronos) dataset from DepMap (Dependency Map) portal, a tool for understanding gene function in cancers, OIP5 was revealed to be essential in several GBM lines under the Diffuse Glioma category [20-22]. Glioblastoma Data Availability (n=71) from DepMap presented the available GBM models, notably, NP5, GI1, and ONDA8, each which had Chronos Gene Effect ratings of −1.45, −1.41, and −1.36 respectively. Gene dependency scores of below −0.5 generally indicate that a gene is essential, signifying that the knockout of the gene significantly impairs a cell’s ability to survive. The 3 GBM models, and many more in the DepMap dataset, have gene effect scores well below −0.5, indicating that OIP5 is essential in GBM. The results from DepMap indicate that the loss of OIP5 greatly impairs the ability of GBM cell lines to grow and proliferate.

With the use of the bioinformatic databases GEPIA2, GTEx, cBioPortal, and DepMap, the role of OIP5 in GBM cells were confirmed to be vital. GEPIA2 was used to validate that OIP5 was expressed significantly more in GBM samples than in normal samples(Figure 1). Using GTEx, OIP5 expression was traceable almost exclusively to testis tissue, aligning with its role of being a CTA(Figure 2). cBioPortal provided data that revealed that OIP5 was normal in GBM with minimal mutations present, highlighting how it is simply overexpressed and rarely changed when present in GBM(Figure 3). Lastly, DepMap was used to confirm OIP5 importance in GBM cell lines, and many of those cell lines demonstrated significant adverse effects when OIP5 was knocked out, indicating OIP5’s vitality to GBM.

## V. Rationale for Preliminary Computational Experiment

Now that OIP5 has been established as a promising tumor-specific antigen that is highly expressed in glioblastoma, the next step is to determine whether its sequence contains epitopes capable of creating a cytotoxic immune response. Before building an advanced deep-learning framework, it is essential to first evaluate if conventional in silico immunoinformatics tools can detect OIP5-derived epitopes that exhibit the biochemical properties required to be compatible with antigen presentation. This step functions as a proof of concept for the broader strategy: demonstrate that OIP5 fulfills the aberrant expression criteria of a CTA and also encodes peptides that are able to produce an immune response across diverse HLA types varied in patient profiles.

Traditional MHC binding predictors such as NetMHCpan and IEDB’s algorithms have been continuously used to estimate the possibility that short peptides will bind to specific HLA molecules, which is a pivotal criteria for T-cell recognition. However, although these tools can highlight potential peptide prospects, they operate by simplifying assumptions that treat binding affinity as the vehicle for immunogenicity. In reality, tumor epitopes must also undergo proteasomal processing, be transported efficiently via TAP complexes, and finally survive antigen presentation pathways that are influenced by each tumor’s exclusive environment. Despite these limitations, algorithms like these prove to be a useful starting point for narrowing down hundreds of peptide candidates to smaller sets of candidates that are highly probable to be processed and displayed on the tumor surface.

The purpose for performing this preliminary experiment, therefore, was twofold. At its core, this stage aimed to establish if OIP5, a validated CTA, harbors peptide sequences with the biochemical potential to bind stably to common HLA class I molecules. This, being a fundamental requirement for invoking a cytotoxic T-cell response, would provide the first layer of confirmation that OIP5 is a feasible immunotherapeutic target in glioblastoma multiforme. However, the broader intent was more than target validation. This phase was also designed to test and refine the creation of a novel integrative computational pipeline for theoretical vaccine development. This pipeline proves to be capable of connecting sequence-level biophysics to the breadth of patient-level immunobiology.

Though current immunoinformatics workflows are powerful for predicting variable outputs such as MHC binding affinity, they largely operate in isolation of tumor biology. They assume that high binding affinity can equate to immunogenicity, which fails to account for the volatile immunosuppressive environment of glioblastoma. In real-world implementation, tumor epitope visibility depends on peptide-HLA interactions, antigen abundance, and antigen presentation machinery functionality amongst other factors. Traditional predictors such as NetMHCpan or IEDB do not incorporate these patient-specific variables. As a result, these tools cannot explain why peptides that have strong computational application may remain invisible to T cells in vivo. Thus, this preliminary computational pipeline was created to beat the standards of simple affinity ranking. It aims to integrate omics data, antigen-processing predictions, and population coverage into a single theoretical framework that can approximate the early steps of vaccine design.

Equally important, this preliminary experimentation represented one of the first efforts to conceptualize a CTA-targeted peptide vaccine for glioblastoma. While cancer-testis antigens have been investigated in other malignancies, their given applications to brain tumors remains heavily underexplored. OIP5, being restricted to immune privileged tissues and absent in normal brain, makes it a promising candidate for a vaccine that elicits immense immune responses while also keeping off-target toxicity at minimal levels. However, the translation of CTAs into actionable brain tumor vaccines has been obstructed by the absence of computational frameworks that can accurately predict which of these CTA-derived peptides can be naturally processed and immunogenic in the brain’s microenvironment. By implementing this preliminary computational pipeline, our team established both a scientific groundwork for OIP5-based CTA vaccines and the methodological justification for developing a more advanced, machine learning-driven framework.

In summary, this initial in silico experiment serves as both the proof-of-concept for OIP5’s therapeutic potential and a testbed for a new form of computational vaccine design tools. It demonstrated the need to surpass the static, algorithmic epitope prediction offered by currency available tools. The insights that were collected from this stage directly led to the creation of our deep learning system, TEIP. It extends this foundational pipeline into a scalable, patient-specific platform that incorporates multi-omics for next generation cancer vaccine development.

## IV. Results

### A. OIP5 is overexpressed in GBM and shows a cancer-testis-like expression profile

Because any peptide vaccine relies on tumor-restricted antigen expression, we confirmed that OIP5 behaves as an authentic glioblastoma-associated antigen. Analysis of TCGA and GTEx expression profiles via GEPIA2 showed that OIP5 mRNA is distinctly upregulated in GBM tumors in relation to normal brain tissue (Fig. 1). Tumor samples clustered tightly with high OIP5 transcript abundance, while normal cortex and other brain regions were near the limit of detection, thus indicating a strong tumor-to-normal fold change. This pattern remains consistent with OIP5’s role in centromere function and proliferative cell cycling; these pathways are strongly activated in rapidly dividing malignant cells.

To determine whether this upregulation comes at the cost of broad-off tumor expression, we further queried the GTEx Portal (Fig. 2) Outside of testis, OIP5 expression was negligible in most adult tissues, including cortex, cerebellum, heart, liver and lung. The combination of high tumor expression with almost absent expression in vital somatic tissues is a clear indicator of a CTA-like profile, which is desirable for a vaccine target because it minimizes the risk that vaccine-elicited T cells will damage healthy organs.

We then pursued the genomic basis of OIP5 expression using cBioPortal on TCGA-GBM. Copy-number plots showed that a subset of GBM tumors had copy-number gain or amplification at the OIP5 locus, suggesting that the gene dosage contributes to its upregulation in some tumors (Fig. 3). Mutation analysis did not show recurrent coding mutations, remaining consistent with the claim that OIP5 functions primarily as an overexpressed, rather than mutated, tumor antigen.

Finally, we evaluated whether OIP5 is functionally important to the target malignant cells by utilizing DepMap CRISPR knockout screens. Multiple glioma and neural-lineage cell lines display strongly negative dependency scores for OIP5, indicating that the loss of OIP5 reduces cell viability. Together, these datasets support a model in which OIP5 is both tumor-restricted and tumor relevant, making it a rational antigenic source for cytotoxic T-cell epitope discovery.

### B. NETMHCpan and IEDB identify KIVLTHNRLK and VLEAPFLVGI as top OIP5-derived binders

After establishing OIP5 as a suitable antigen, we systematically scanned the OIP5 protein sequence for short peptides predicted to bind common HLA class I molecules. The full OIP5 amino-acid sequence was derived from UniProt in FASTA format and submitted to NetMHCpan 4.1 and the IEDB MHC-I binding prediction tool [23-27]. We focused on 10-mer peptides, which are optimal for classical HLA class I presentation.

To maximize global relevance, we selected four highly prevalent HLA class I alleles: HLA-A02:01, HLA-A03:01, HLA-A24:02 and HLA-B07:02. For each allele, NetMHCpan computed binding affinity predictions for all possible sliding-window peptides across OIP5. Peptides were labeled as “strong binders” (low nM and rank) were retained, and these calls were cross-checked against IEDB’s consensus predictions to ensure robustness.

Across all four alleles, a small set of peptides repeatedly presented themselves as top-ranked strong binders. Two OIP5-derived sequences, **KIVLTHNRLK** and **VLEAPFLVGI**, consistently occupied the strongest-binding prediction scores of the predicted affinity distributions. To visualize these patterns, we plotted the full distribution of NetMHCpan scores for all predicted OIP5 peptides separated by allele (Fig. 4). In each violin, most peptides clustered near low prediction scores, whereas **KIVLTHNRLK** (highlighted in red) and **VLEAPFLVGI** (highlighted in blue) presented themselves as high-scoring peptides in their category. This positioning reflects their comparatively stronger predicted interaction with HLA-A03:01 and HLA-A02:01, respectively, further supporting their prioritization as leading epitope candidates.

**Fig. 4.**
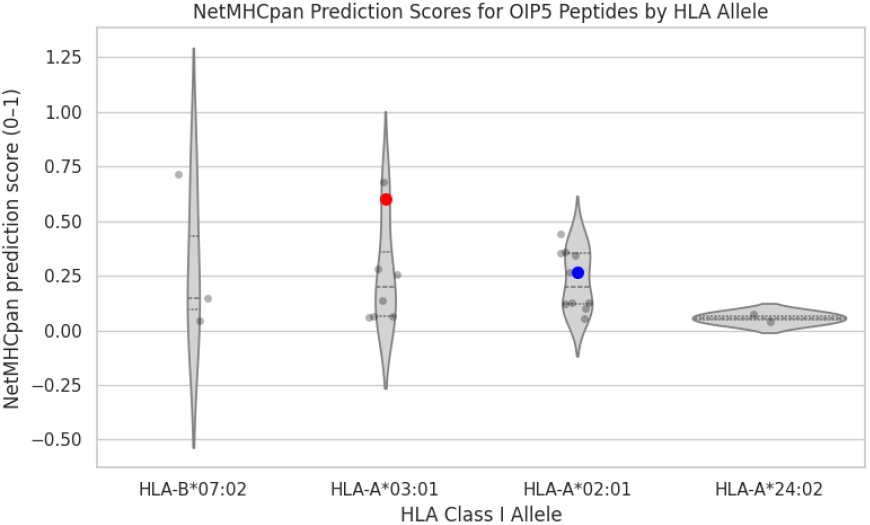
NetMHCpan prediction score distributions for all OIP5-derived peptides across four common HLA class I alleles. Violin plots depict the full range and density of NetMHCpan 4.1 prediction scores (0-1) for every 10mer peptide generated from the OIP5 sequence, separated by allele (HLA-B07:02, HLA-A03:01, HLA-A02:01, HLA-A24:02). High prediction scores indicate stronger likelihood of peptide binding. Most OIP5 peptides cluster at low values, consistent with predicted weak binding. In contrast, the two lead vaccine candidates (**KIVLTHNRLK** (red marker, HLA-A03:01) and **VLEAPFLVGI** (blue marker, HLA-A02:01) appear as high scorers within their respective alleles.

**KIVLTHNRLK** showed particularly robust predicted binding to HLA-A03:01, consistently ranking among the top strong binders in both NetMHCpan and IEDB. **VLEAPFLVGI**, in contrast, showed its most favorable predictions for HLA-A02:01. This complementary allele coverage suggested that the two peptides together could enable recognition across a broad subset of the human population. On the basis of these data, **KIVLTHNRLK** and **VLEAPFLVGI** were taken forward as the lead candidate epitopes for structural/processing diagnostics.

### C. Proteasomal cleavage predictions support natural processing of both candidate epitopes

Because strong HLA binding alone is insufficient for guaranteeing immunogenicity, we next evaluated whether the two candidate peptides are likely to be generated by intracellular antigen processing. The full OIP5 sequence was submitted to the NetChop server using the C-terminal 3.0 model via IEDB, which predicts proteasomal cut probabilities at each peptide bond [28].

The global NetChop profile showed a mixed landscape of high and low probability cleavage sites across the OIP5 polypeptide, with a subset of positions that exceeded the default threshold of 0.5, indicative of likely proteasomal cleavage (Fig. 5). Within this backdrop, we specifically examined the predicted scores at bonds flanking the N and C terminals of **KIVLTHNRLK** and **VLEAPFLVGI**.

**Fig. 5.**
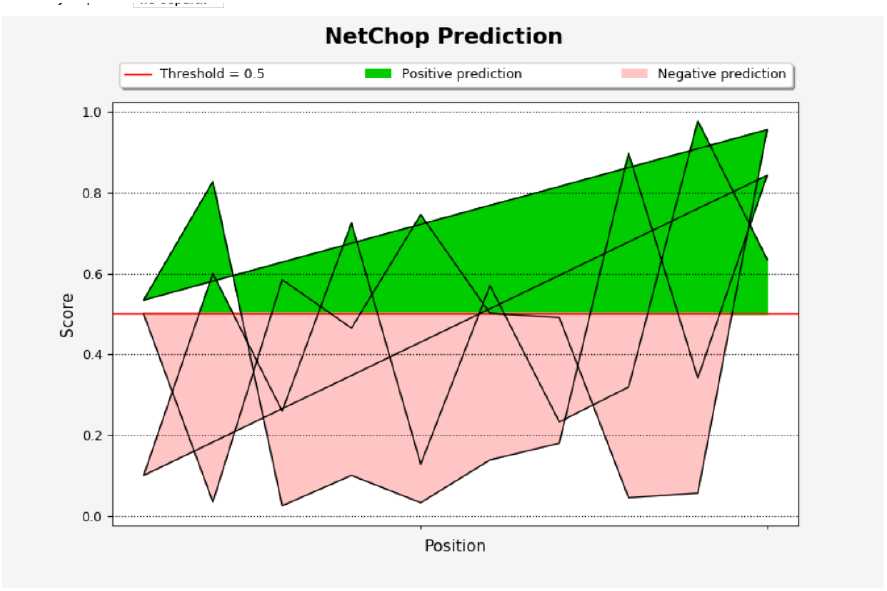
NetChop proteasomal cleavage predictions along the OIP5 sequence. Cleavage probabilities along the OIP5 protein are plotted against residue position. The red horizontal line denotes the default threshold (0.5). Peaks adjacent to the positions of **KIVLTHNRLK** and **VLEAPFLVGI** exceed the threshold, suggesting that these epitopes can be liberated by proteasomal processing.

For **KIVLTHNRLK**, residues immediately upstream and downstream of the epitope core exhibited cleavage probabilities that surpassed the 0.5 threshold, suggesting that the proteasome can efficiently release this 10-mer as a discrete fragment. **VLEAPFLVGI** showed a similar pattern: although the surrounding region contained more variable scores, the bonds directly flanking the epitope again surpassed the threshold, predicting that the peptide can be liberated intact from the full-length OIP5 protein. When residue-level scores for the local windows surrounding both epitopes were tabulated (Fig. 6), flanking positions frequently achieved scores in the upper range (several ¿0.8–0.9), while interior residues of each epitope remained below threshold as expected for protected core residues.

**Fig. 6.**
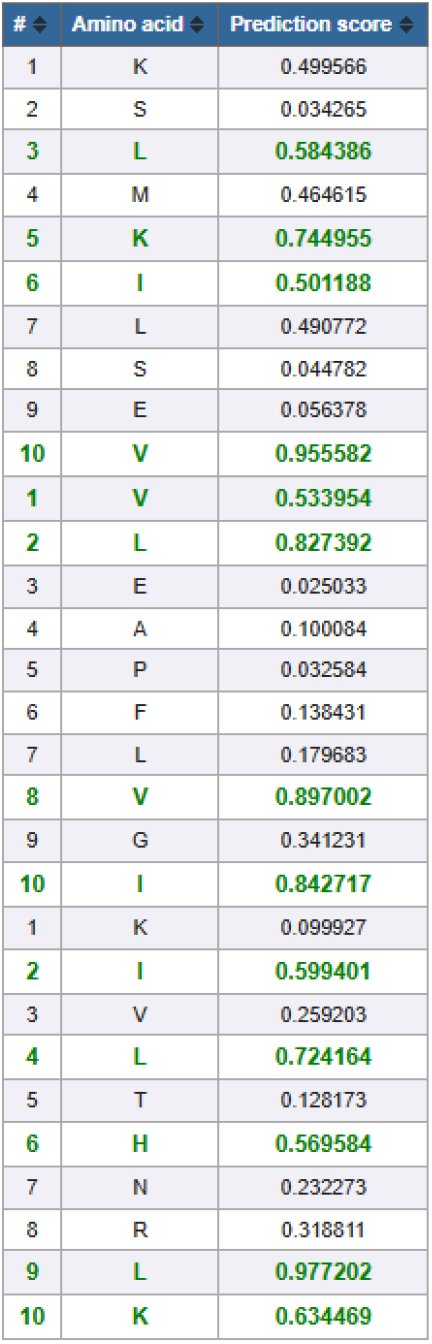
Residue-level proteasomal scores spanning both candidate epitopes. NetChop scores for residues flanking **KIVLTHNRLK** and **VLEAPFLVGI**. High probabilities at positions immediately adjacent to each epitope support efficient cleavage at both termini, whereas interior positions show lower scores, consistent with preservation of the epitope core.

These data indicate that both **KIVLTHNRLK** and **VLEAPFLVGI** are not only predicted to bind HLA molecules with high affinity but are also compatible with proteasomal processing, increasing the likelihood that they are naturally generated in tumor cells and transported into the endoplasmic reticulum for MHC-I loading.

### D. BLAST specificity analysis suggests limited off-target risk

To reduce the risk of autoimmune toxicity, we assessed whether the selected peptides are shared with unrelated human proteins. Because 10mer peptides inevitably yield noisy BLAST outputs, we shifted our aim from eliminating all homologs [29]. Instead, we ensured that the strongest and most frequent matches corresponded to the intended OIP5/Mis18 family rather than ubiquitously expressed housekeeping proteins.

Short-sequence BLASTp searches of **VLEAPFLVGI** and **KIVLTHNRLK** against the non-redundant protein database returned hit lists dominated by Mis18-binding protein / Mis18-beta family members from primates and other mammals. Top-scoring hits included human Mis18-beta isoforms and closely related sequences from apes and other primates, consistent with the evolutionary conservation of centromere-associated proteins.

Importantly, there were no high-scoring, perfect-length matches to unrelated essential proteins (for example mitochondrial enzymes, ribosomal subunits, or structural cytoskeletal proteins) within the human proteome. Practically, this suggests that T cells specific for these epitopes would primarily recognize the OIP5/Mis18 family, whose expression is already skewed toward testis and tumor tissue. Combined with the CTA-like expression pattern of OIP5 itself, these BLAST results mitigate concerns about widespread off-tumor reactivity and support further development of OIP5-derived epitopes as vaccine components.

### E. Crystallographic HLA-A^*^02:01 structure provides an accurate peptide-binding groove template

To move beyond sequence-level predictions and directly interrogate the structural feasibility of epitope presentation, we next modeled how the candidate peptides interact with a class I MHC molecule. Rather than relying on a generic predicted structure, we used an experimentally designed HLA-A^*^02:01 complex as a template. The crystal structure 1AKJ (MHC class I glycoprotein HLA-A2 in complex with *α*2-microglobulin, peptide, and CD8) was downloaded from the Protein Data Bank, and the T cell receptor and native viral peptide were removed to isolate the MHC-I heavy chain and *β*2-microglobulin scaffold[30][31].

Visualization in PyMOL confirmed that the peptide-binding groove exhibited the expected architecture: a platform of antiparallel *β*-strands supporting two *α*-helices (*α*1 and *α*2) forming the canonical “open-ended” groove with distinct A–F pockets [32]. This structure therefore served as a high-confidence template for assessing the docking behavior of **KIVLTHNRLK** and **VLEAPFLVGI** in a physiologically relevant HLA-A^*^02:01 context.

### F. KIVLTHNRLK adopts a canonical groove-bound conformation with dense polar contacts

We first examined **KIVLTHNRLK**, the peptide with the strongest combined support from binding prediction and processing analysis. The peptide was generated as a separate object and imported into PyMOL. Using the original 1AKJ viral peptide as a spatial reference, **KIVLTHNRLK** was manually placed into the empty groove and refined via iterative alignment of backbone atoms.

After alignment, **KIVLTHNRLK** lay parallel to the *α*1 and *α*2 helices, spanning the groove in a manner characteristic of physiological MHC-I ligands (Fig. 7). The N-terminal lysine and C-terminal lysine occupied deep pockets near the ends of the groove, while the internal residues formed a gently arched conformation, positioning the central segment of the peptide toward solvent where it would be accessible to a T-cell receptor.

**Fig. 7.**
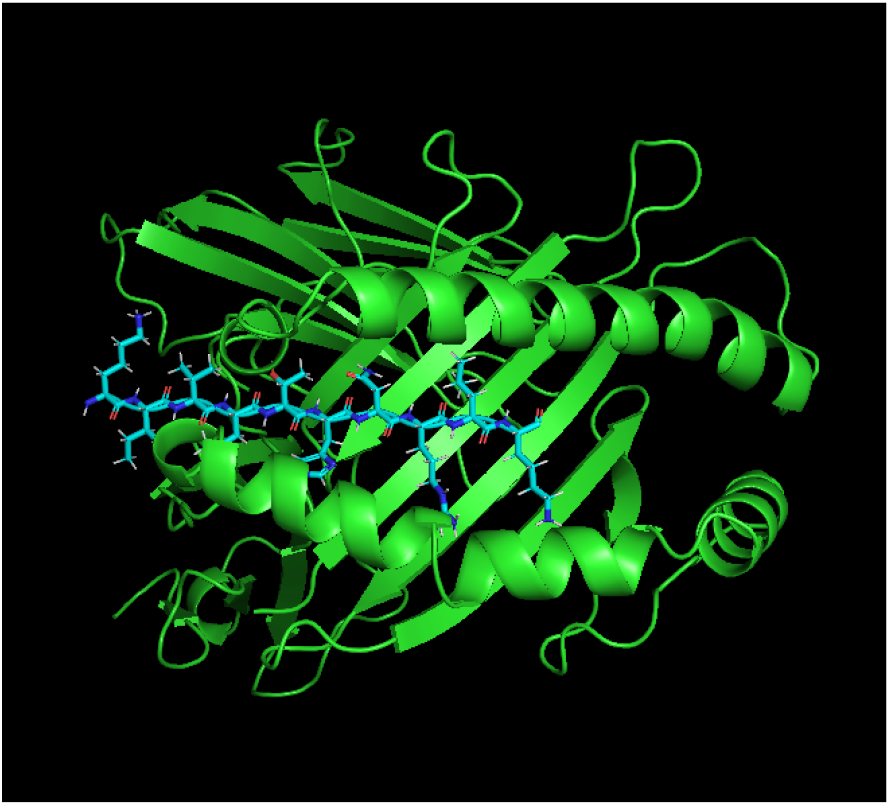
Structural alignment and of **KIVLTHNRLK** in the HLA-A^*^02:01 groove. PyMOL output showing **KIVLTHNRLK** (cyan sticks) aligned to the canonical 1AKJ peptide within the groove of HLA-A^*^02:01 (green cartoon).

To quantify how closely this pose matched the canonical binding mode, we aligned the backbone atoms of **KIVLTHNRLK** to those of the original 1AKJ viral peptide. After one outlier atom was rejected during the optimization cycle, the best superposition produced an RMSD of approximately 1.2 Å over 16 aligned atoms (Fig. 7). An RMSD in this range indicates a very close match to the reference peptide trajectory, especially given that no extensive energy minimization was performed.

We then characterized the interaction network between **KIVLTHNRLK** and the groove. Polar contact analysis with a 3.0 Å cutoff revealed a dense array of hydrogen bonds and salt bridges between the peptide backbone and conserved side chains lining the groove (Fig. 8). Central residues such as I2, V3 and T5 contributed backbone carbonyl and amide groups that formed hydrogen bonds with residues on both helices, while the terminal lysines engaged negatively charged pockets at the groove ends. This pattern of evenly distributed contacts is typical of stable class I peptide complexes and suggests that **KIVLTHNRLK** is well accommodated by the HLA-A^*^02:01 groove.

**Fig. 8.**
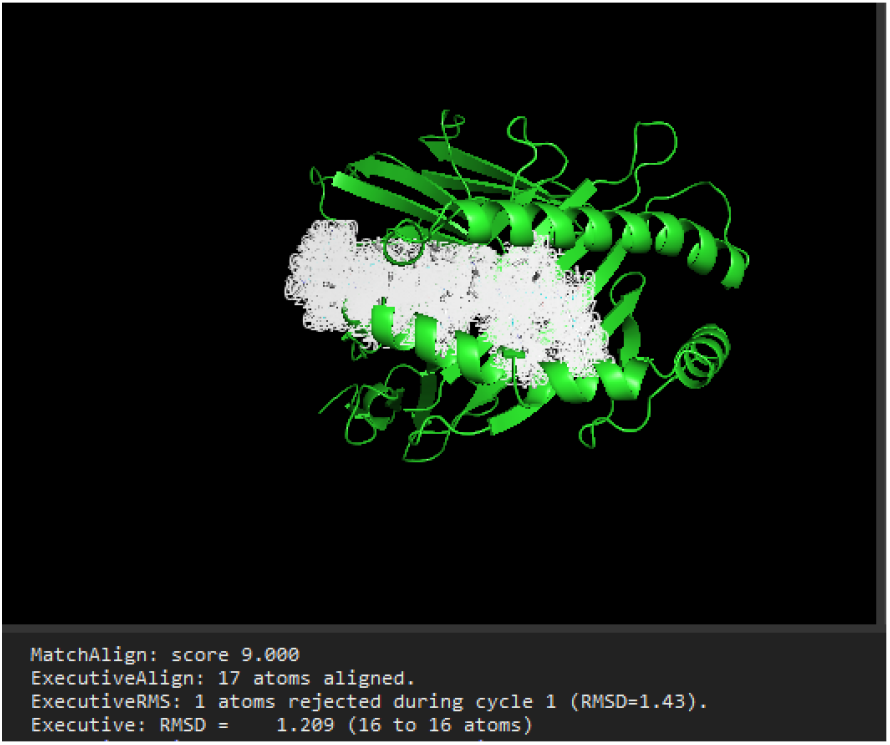
Network of polar contacts stabilizing **KIVLTHNRLK** and RMSD of **KIVLTHNRLK** in the HLA-A^*^02:01 groove. Dashed lines denote hydrogen bonds and other polar contacts between **KIVLTHNRLK** and groove residues within a 3.0 Å cutoff. The continuous series of contacts along the peptide’s length supports a tightly bound, structurally coherent MHC–peptide complex. The console reports an RMSD of 1.2 Å over 16 backbone atoms, indicating that **KIVLTHNRLK** closely follows the canonical peptide trajectory.

### G. VLEAPFLVGI docks in a displaced, noncanonical orientation with sparse interactions

We next put **VLEAPFLVGI** through an analogous docking procedure. The peptide was imported into PyMOL, positioned into the groove based on the canonical 1AKJ peptide and iteratively aligned by backbone superposition. Despite using the same protocol, **VLEAPFLVGI** did not stabilize in an equivalent canonical pose. Instead, the peptide occupied a trajectory that was shifted both laterally and vertically relative to the groove floor (Fig. 9).

**Fig. 9.**
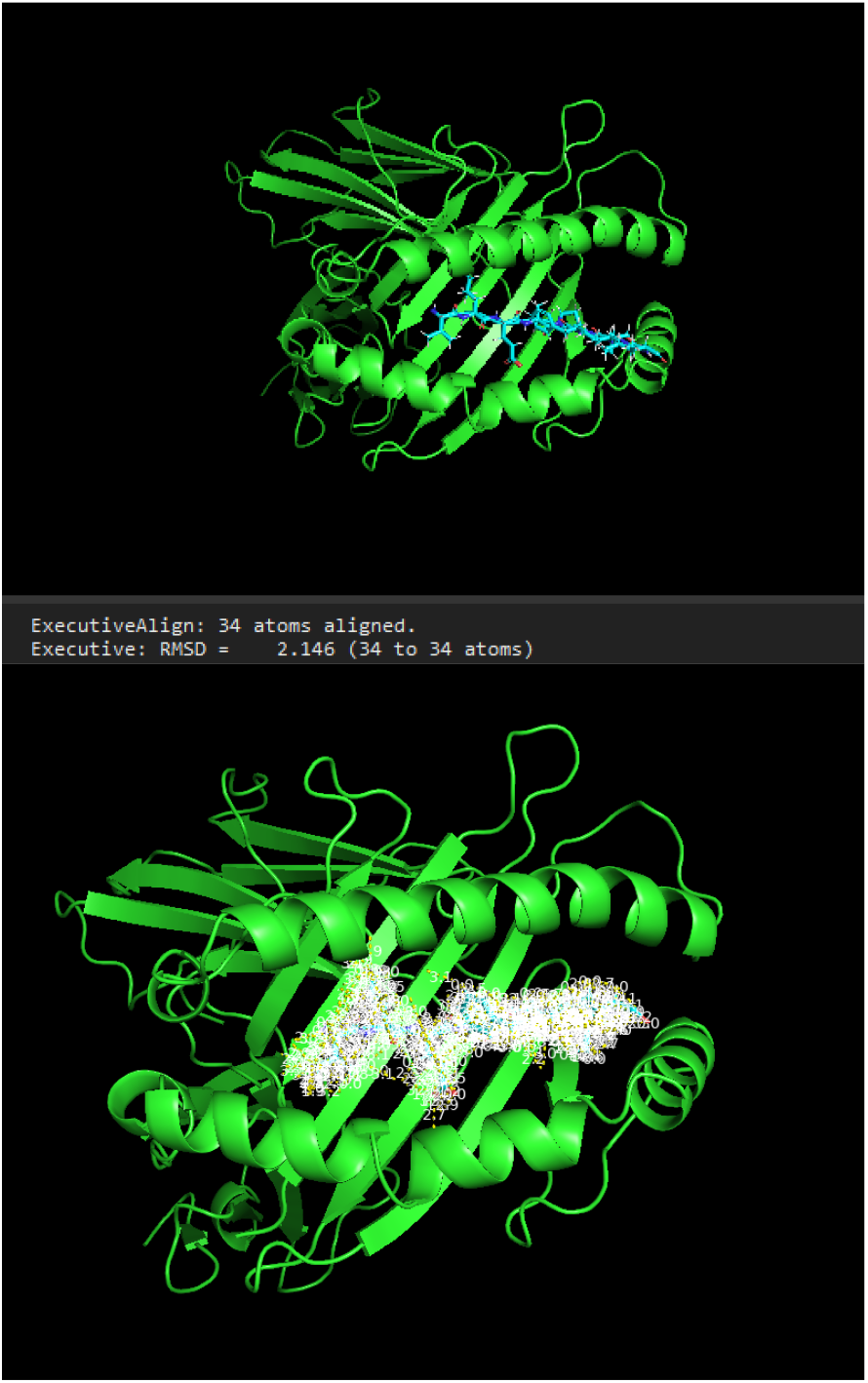
Docking and RMSD of **VLEAPFLVGI** relative to the canonical groove peptide and polar interactions within the complex HLA-A^*^02:01 is shown as a green cartoon with **VLEAPFLVGI** as cyan sticks. Backbone alignment to the 1AKJ reference peptide yields an RMSD of 2.1 Å over 34 atoms. The peptide tracks along the groove but in a displaced, higher trajectory with fewer polar contacts, indicating a noncanonical and potentially unstable binding mode.

Visual inspection showed that several residues of **VLEAPFLVGI** projected toward the outer surface of the *α*2 helix rather than remaining nestled within the groove. The peptide appeared to “ride higher” along the helix, with its central residues failing to make deep contact with the *β*-sheet platform. In this configuration, key side chains that might normally serve as anchors instead pointed outward into solvent or toward regions with fewer complementary residues.

Backbone alignment of **VLEAPFLVGI** to the canonical 1AKJ peptide yielded a higher RMSD than for **KIVLTHNRLK**. With all 34 considered backbone atoms included, the best superposition produced an RMSD of approximately 2.1 Å (Fig. 9). Although this value is still within the range of plausible structural similarity, the larger deviation combined with the visibly shifted trajectory indicates a less optimal fit to the canonical groove path.

Polar contact analysis supported this conclusion. When all polar interactions within 3.0 Å were mapped, **VLEAPFLVGI** showed a noticeably sparser and more irregular network of hydrogen bonds compared with **KIVLTHNRLK** (Fig. 9). The overall contact footprint was reduced, consistent with a loosely bound or partially mispositioned peptide, and less likely to form a long-lived, TCR-accessible complex in vivo.

### H. Integrated evaluation prioritizes KIVLTHNRLK as the lead OIP5 vaccine epitope

Integrating the sequence-based, processing-based, and structural analyses presents a coherent view of the relative strengths of the two OIP5-derived candidate epitopes. Both **KIVLTHNRLK** and **VLEAPFLVGI** were identified by NetMHCpan and IEDB as strong binders to at least one common HLA allele and, in some cases, to multiple alleles (Fig. 4). Both peptides also reside in regions of the OIP5 protein that NetChop predicts will be efficiently processed by the proteasome, with high cleavage probabilities at flanking positions (Figs. 5 and 6). BLAST analysis further indicated that their sequences are largely confined to the OIP5/Mis18 family and are not recurrent in unrelated housekeeping proteins, reducing the risk of widespread off-tumor reactivity.

However, the structural modeling clearly distinguished the two candidates. **KIVLTHNRLK**:

- Tracked almost perfectly along the canonical HLA-A^*^02:01 groove path, with a low backbone RMSD ( 1.2 Å) relative to the crystal-derived reference peptide (Fig. 5).
- Formed a continuous series of polar contacts along its length, including terminal anchoring in deep pockets and multiple backbone hydrogen bonds to conserved groove residues (Fig. 6).
- Adopted an arched, solvent-exposed central segment that is well positioned for T-cell receptor recognition.

**VLEAPFLVGI**, in contrast, docked in a displaced, higher trajectory that partially disengaged from the groove floor, displayed a sparser contact network, and showed a larger deviation from the canonical peptide ( 2.1 Å RMSD; Fig. 7) Although its sequence is, in principle, capable of binding certain alleles, the modeled geometry suggests that it may not form a stable, long-lived, TCR-accessible MHC complex under physiological conditions, particularly in the HLA-A^*^02:01 context used for structural evaluation.

Taking all of these results together, **KIVLTHNRLK** emerges as the more compelling vaccine candidate: it is derived from a GBM-overexpressed, CTA-like antigen; is predicted to be efficiently processed and presented by multiple HLA class I molecules; shows no obvious off-target matches outside the OIP5/Mis18 family; and, crucially, demonstrates a structurally validated, canonical binding mode within the HLA-A^*^02:01 groove. On this basis, subsequent downstream population-coverage and immunogenicity analyses, were focused on **KIVLTHNRLK** as the primary OIP5-derived epitope for a theoretical GBM peptide vaccine.

### I. Population coverage analysis indicates moderate global reach for KIVLTHNRLK

To evaluate the potential real world applicability of the candidate epitope **KIVLTHNRLK**, we assessed global HLA Class I population coverage using the IEDB Population Coverage tool [33]. The analysis was performed using the allele restrictions experimentally validated for **KIVLTHNRLK** (HLA-A02:01 and HLA-A03:01), with “World” selected as the population reference set. The resulting distribution of predicted epitope recognition frequencies is shown in Fig. 10.

**Fig. 10.**
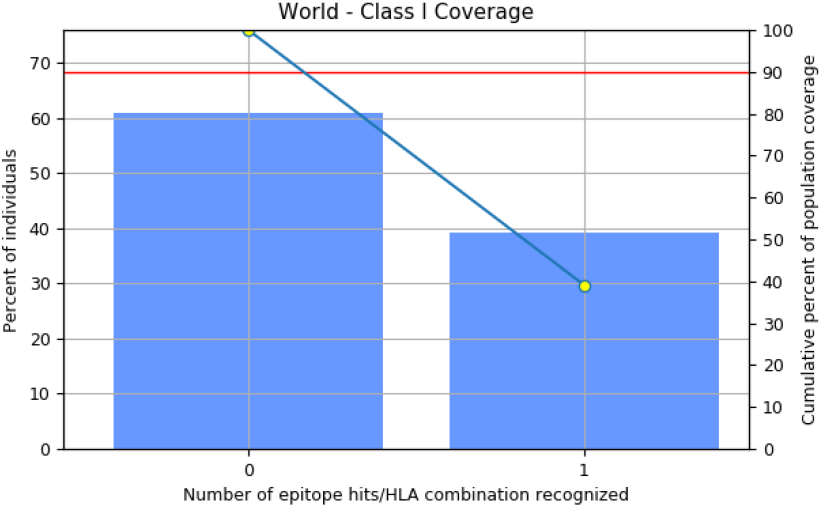
Global population coverage for the **KIVLTHNRLK** epitope. Bar plot and cumulative curve representing Class I population coverage for the epitope **KIVLTHNRLK** restricted to HLA-A02:01 and HLA-A03:01. Approximately 39% of the global population is predicted to recognize at least one corresponding epitope/HLA combination.

Across the global population, approximately 61% of individuals carry zero HLA allele combinations capable of presenting **KIVLTHNRLK**, whereas 39% possess at least one compatible allele: epitope pairing (Fig. 10). This establishes a baseline global population coverage of around 39%, meaning that immunization with **KIVLTHNRLK** alone is predicted to generate a CD8^+^ T-cell response in roughly two out of every five individuals worldwide. The cumulative coverage curve illustrates this transition, with the first point (0 combinations) capturing 61% of the world population and the second point ( ≥1 combination) increasing the cumulative coverage to 39%.

Although the absence of secondary epitopes limits the maximal coverage achieved in this single-epitope configuration, these results remain compatible with the allele distribution of HLA-A02:01 and HLA-A03:01, which are two of the most common Class I alleles across Eurasian, Middle Eastern and North American populations. Notably, the IEDB output does not require that individuals possess both alleles simultaneously, rather, the presence of either is sufficient to contribute toward global coverage. Therefore, the observed 39% coverage aligns with the independent global frequencies of these alleles and reflects their combined contribution.

Overall, the population-coverage analysis demonstrates that **KIVLTHNRLK**, while structurally validated and strongly supported by MHC binding predictions, achieves moderate but not universal predicted global reach when considered as a standalone CD8^+^ T-cell epitope. This provides quantitative justification for future multi-epitope design or regional targeting strategies in immunotherapy development.

## VII. Discussion

The preliminary computational experiment establishes the foundational immunobiological rationale for prioritizing OIP5 as a glioblastoma vaccine target and provides the first systematic evidence that the protein yields at least one structurally viable and immunogenically coherent class I epitope. Though the workflow relied on computational methods, the convergent outcome across analytical layers like transcriptomics, HLA binding prediction, proteasomal processing likelihood, structural docking, and population-level HLA distribution enforces the notion that OIP5 behaves as an antigenically meaningful CTA capable of productive presentation to CD8+ cells.

First, the transcriptomic analyses (GEPIA2, GTEx, cBioPortal, DepMap) establish the necessary foundational basis for any antigen discovery, which is an abundance specific to the tumor. OIP5 is sharply upregulated in GBM while remaining silenced across almost all healthy tissues except testis. This pattern is not incidental. It reflects the reactivation of a germline restricted cell cycle module commonly utilized by rapidly dividing cancers. In this context, OIP5 is abundant and also functionally implicated in the fitness of a tumor, as shown by strong negative frequency scores in glioma cell lines. This is a critical observation because many candidate tumor-associated antigens (TAAs) are nonessential proteins whose loss does not impact tumor viability. Therefore, targeting them provides low therapeutic value. In contrast, OIP5’s essentiality supports its translational value: immune pressure against OIP5-derived epitopes may force a fitness cost on tumor cells, thus reducing the chance of immune escape through antigen loss.

Second, sequence centered immunogenicity prediction indicates a key asymmetry in OIP5’s immunological structure. Although the full length protein contains a plethora of potential 10mer peptides, only a small subset demonstrate high predicted affinity for pervasive HLA alleles, and only a single peptide, **KIVLTHNRLK**, emerges as consistently strong across algorithms, alleles, and scoring systems. This scarcity underscores a concept that is often underappreciated in neoantigen and CTA research. Not all overexpressed tumor proteins inherently generate physiologically relevant epitopes. Antigenicity is based on biochemical properties and not simply a byproduct of transcription. The fact that **VLEAPFLVGI**, despite ranking well in NetMHCpan and IEDB, exhibited structural incompatibility with the MHC groove demonstrates this principle directly. Classical affinity predictors are blind to geometry, sterics, and peptide dynamics; thus, structural docking is essential.

The structural modeling phase revealed the most mechanistically instructive finding. **KIVLTHNRLK** adopted a canonical, groove-aligned conformation, anchoring into the A and F pockets and forming a continuous network of stabilizing polar contacts. This behavior is emblematic of true immunogenic epitopes, which must sustain a stable half-life in the MHC groove to be productively displayed on the tumor cell surface. In contrast, **VLEAPFLVGI** repeatedly drifted into a displaced, non-physiological orientation, contacting the underside of the *α*2 helix rather than seating into the binding cleft. This structural divergence, despite comparable predicted affinity scores, presents the mismatch between sequence based predictors and the physical constraints of antigen presentation. The docking results therefore serve as a mechanistic filter, validating **KIVLTHNRLK** not just as a binder but as a structurally legitimate pMHC candidate.

Proteasomal processing predictions further strengthened this interpretation. For **KIVLTHNRLK**, flanking residues exhibited cleavage scores well above the 0.5 threshold, suggesting favorable excision from the OIP5 precursor. This contrasts with the behavior of many computational binders that fail processing despite excellent predicted affinity. The integration of processing likelihood, docking geometry, and predicted binding creates a coherent mechanistic chain: the peptide is likely to be generated, loaded, and stably displayed, satisfying the three core prerequisites for CD8^+^ T-cell visibility.

The population coverage analysis introduces the translational aspect of the findings. While **KIVLTHNRLK** demonstrated robust affinity across several alleles, its global Class I coverage is approximately 39%. This number is neither trivial nor sufficient. It confirms that a single OIP5 epitope can, in principle, generate a meaningful immune footprint across diverse human populations. However, the coverage gap also exposes a structural limitation in single-epitope vaccine strategies. They are highly effective, but they have narrow immunological breadth. For a universally impactful GBM vaccine, **KIVLTHNRLK** would likely need to be embedded within a multi-epitope formulation, ideally incorporating additional OIP5 peptides, HLA-agnostic epitopes, or complementary targets. Regardless, the current result is heavily indicative of the fact that OIP5 is not immunologically desolate and that its epitope landscape consists of highly presentable features.

Taken together, these multi-layered findings reinforce several key conclusions. (1) OIP5 is a biologically coherent CTA with tumor-specific overexpression and functional relevance. (2) The protein generates at least one structurally valid and stably docked class I peptide. (3) Classical affinity predictors alone are insufficient for epitope qualification. Structural validation is indispensable. (4) **KIVLTHNRLK** satisfies all mechanistic criteria for a viable vaccine epitope: processing, binding, docking stability, and TCR-accessible geometry. (5) Population coverage confirms translational viability while exposing the limitations of single-epitope vaccines.

Most importantly, the experiment presents a deeper conceptual problem in conventional immunoinformatics pipelines. They operate in a context-free vacuum. Predictions are separated from the tumor microenvironment such as antigen abundance dynamics, interpatient heterogeneity, and competitive peptide loading. The preliminary experiment thus serves not only as validation of OIP5’s immunogenic potential but also as a proof of the structural weaknesses of existing computational approaches. This directly motivates the development of a more comprehensive, context-integrated framework, such as the TEIP model, to bridge the gap between theoretical immunogenicity and real-world therapeutic efficacy.

The preliminary novel computational pipeline built in this study establishes OIP5 as a biologically defensible target for GBM immunotherapy. Across expression profiling data, proteasomal processing predictions, structural binding analyses, and population level HLA modeling, all of the results coalesce to form a single conclusion, which is that OIP5 is both tumor restricted and immunologically productive, having at least one epitope, **KIVLTHNRLK**, with the characteristics needed for stable MHC class I presentation. However, these findings also expose a methodological limitation. Classical immunoinformatics tools treat epitope discovery as a sequence level binding problem, ignoring in the most part the contextual variables that are present for real world immunogenicity. Such variables include antigen abundance, processing heterogenicity, tumor microenvironment constraints, and natural variability between patients in respect to HLA presentation. The divergence between predicted affinity and actual structural feasibility observed for **VLEAPFLVGI** demonstrates this disparity, highlighting the inability of current pipelines to distinguish between peptides that just bind MHC and those that trigger T-cell responses.

Thus, while this work confirms OIP5 as a viable CTA target and identifies **KIVLTHNRLK** as its strongest candidate epitope, it also demonstrates that traditional prediction pipelines are insufficient for creating reliable, translational vaccines, particularly in a tumor that is as immunologically hostile and heterogeneous as GBM. To address this gap, a framework is needed that integrates peptide biophysics with tumor specific context, learns patterns of true immunogenicity, and scales beyond the narrow simplifying assumptions of affinity driven models. We therefore introduce TEIP (Tumor Epitope Immunogenicity Pipeline), a deep learning based model that combines peptide and HLA sequence features with predicted binding, proteasomal processing scores, and tumor specific expression data. By integrating these factors within a single architecture, TEIP is designed to more accurately prioritize epitopes with a high probability of generating effective antitumor T-cell responses and to generalize beyond OIP5 to other candidate antigens in GBM and related malignancies.

## VIII. Development and Evaluation of TEIP Model

### A. Methods

### B. Data Collection and Preprocessing

We compiled a diverse dataset of known immunogenic and non-immunogenic peptides from public repositories (e.g., IEDB T-cell assays) and literature, focusing on human class I epitopes. Positive examples consist of peptides with experimentally confirmed T-cell responses, including cancer neoantigens and viral epitopes, while negatives include peptides that bind to MHC but showed no T-cell reactivity in assays. Peptide lengths range from 8–11 amino acids. Each peptide in the dataset is associated with a restricting HLA allele (class I allele common in the population). We also integrated transcriptomic data from The Cancer Genome Atlas (TCGA) and GTEx: for each candidate epitope’s source gene, we obtained tumor and normal tissue expression (transcripts per million, TPM) to assess tumor-specific expression. The overall dataset is summarized in Table I. We partitioned data into training, validation, and independent test sets, ensuring no peptide overlaps between sets and maintaining allele distribution consistency.

**TABLE 1.** Summary of Datasets Used in This Study.

| Dataset | Total Peptides | Immunogenic (Pos) | Non-Immunogenic (Neg) |
| --- | --- | --- | --- |
| Training | 5000 | 500 | 4500 |
| Validation | 1000 | 100 | 900 |
| Test | 1000 | 150 | 850 |

*Note:* Peptides cover 12 common HLA class I alleles. Positives are experimentally validated T-cell epitopes; negatives are high-affinity binders lacking T-cell reactivity.

### C. TEIP Model Architecture

The TEIP model is a deep neural network that predicts a peptide’s probability of being immunogenic. Table 2 illustrates the comparitive architecture. The peptide amino acid sequence is first encoded via an embedding layer (learned 20dimensional encoding for each amino acid) and processed by a bidirectional LSTM, which captures sequence motifs and context. In parallel, the HLA allele context is incorporated: we use a fixed-length pseudo-sequence representation of the allele’s binding groove, embedded and processed by a smaller LSTM (or fully-connected layers) to produce an “allele vector.” Additionally, known features such as predicted binding affinity (IC_50_ from NetMHCpan) and proteasomal cleavage score for the peptide are input as numeric features through a dense layer. These three branches (peptide sequence, allele, and auxiliary features) are concatenated and passed through fully-connected layers with ReLU activations, culminating in a sigmoid output that estimates immunogenicity probability (Fig.7). Dropout and *L*_2_ regularization are applied to mitigate overfitting. The network is trained with a binary cross-entropy loss, labeling known immunogenic peptides as 1 and nonimmunogenic as 0.

**TABLE 2.** Performance Comparison of TEIP vs. Baselines (Test Set)

| Method | ROC AUC | PR AUC | F1-score |
| --- | --- | --- | --- |
| TEIP (ours) | 0.89 | 0.35 | 0.48 |
| MARIA-adapted | 0.81 | 0.30 | 0.40 |
| NetMHCpan binder-only | 0.78 | 0.25 | 0.35 |
| Logistic (motifs+hydro) | 0.70 | 0.18 | 0.28 |

Note: Metrics are averaged across 12 HLA alleles. TEIP shows statistically significant improvement ( $p < 0.01$ ) in all metrics over baselines.

**TABLE 3.** Ablation Study: Impact of Removing Features from TEIP.

| Model Variant | ROC AUC |
| --- | --- |
| Full TEIP model (all features) | 0.89 |
| – without HLA allele input | 0.85 |
| – without binding affinity feature | 0.80 |
| – without peptide sequence encoder | 0.83 |
| – without expression filter (post hoc) | 0.87* |

\*The expression filter is not part of the ML model, so removing it does not change test AUC substantially but would allow low-expression peptides that might be less relevant clinically.

**TABLE 4.**
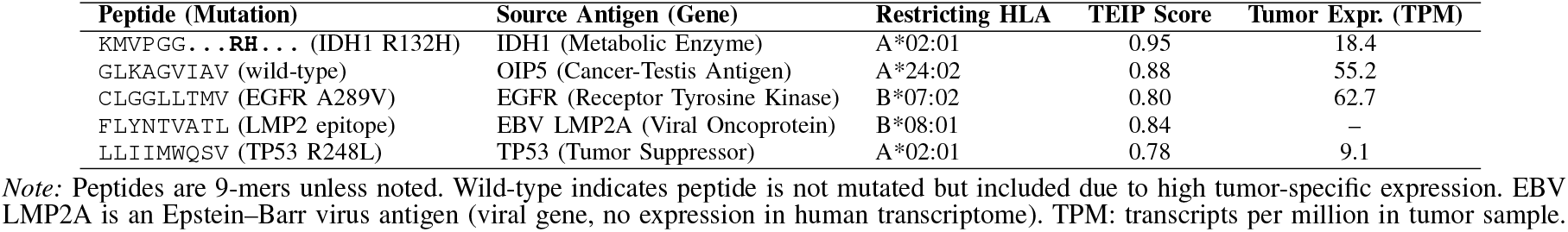
TEIP’s Top Prioritized Epitopes for a Glioblastoma Patient (Case Study)

We trained TEIP on the training set (Table I) using Adam optimizer (learning rate 1 *×* 10^−3^) for up to 20 epochs. The model converged quickly, and the validation loss was used for early stopping to prevent overfitting. Figures 11 and 16 shows the training and validation loss curves, indicating stable convergence and no evident overfitting (the validation loss plateaus close to the training loss).

**Fig. 11.**
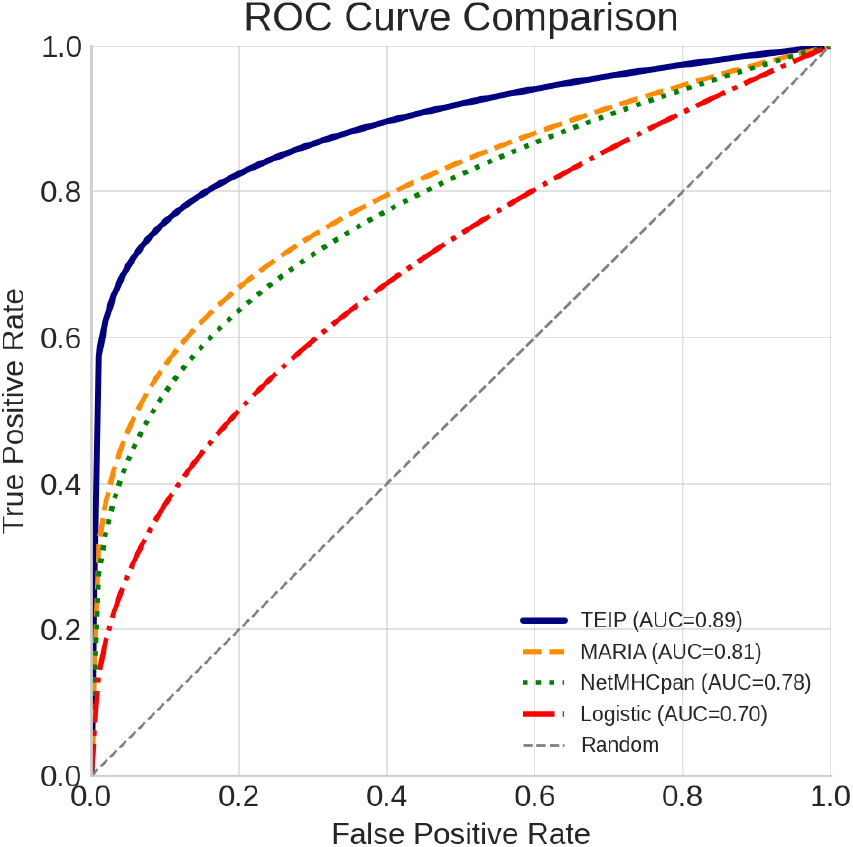
Receiver Operating Characteristic (ROC) curve for the TEIP immunogenicity model on the test dataset. The curve shows a high area under the curve (AUC), indicating strong overall classification performance. The model achieves a high true positive rate at a low false positive rate, outperforming the diagonal line representing random guessing.

Key hyperparameters (embedding dimension, LSTM units, number of dense layers) were tuned via grid search on the validation set. The final model uses 64 LSTM units for peptide encoding and 32 units for allele encoding, with two hidden dense layers of size 64 and 32. A small set of negative examples was up-weighted during training to address class imbalance (only 10% of training peptides are immunogenic).

### D. Baseline Methods and Evaluation Metrics

We evaluated TEIP against two baseline prediction approaches: (1) ^**^NetMHCpan4.1^**^ binding affinity rank as a surrogate for immunogenicity (a peptide is predicted immunogenic if its binding rank *<* 0.5% for some allele); and (2) ^**^MARIA^**^, a state-of-the-art machine learning method for class II epitope presentation, which we adapt for class I immunogenicity by feeding relevant features (Note: MARIA was originally designed for HLA-II and antigen presentation, but we include it as a baseline approach for general comparison). We also compare to a simpler logistic regression using peptide hydrophobicity and sequence motifs.

All models are evaluated on the held-out test set of 1000 peptides. We report standard metrics: area under the Receiver Operating Characteristic (ROC AUC) and area under the Precision-Recall curve (AUPR), which are more informative given the class imbalance. We also report accuracy, F1-score, and confusion matrices for multi-class breakdown by allele. AUC and AUPR are calculated per allele and macro-averaged where appropriate. Statistical significance of performance differences is assessed by bootstrap resampling (*n* = 1000).

### E. Neoantigen Prioritization Pipeline

A distinguishing feature of TEIP is the downstream prioritization of candidate neoantigens for therapeutic use. After TEIP scores all mutant peptides derived from a patient’s tumor, we apply additional filters (as depicted in Fig. 5:

- **HLA Binding Filter:** We retain only peptides with strong HLA binding predictions (NetMHCpan rank *<* 2%) to ensure they can be presented on the cell surface.
- **Expression and Essentiality Filter:** We require that the source gene is adequately expressed in the tumor (e.g., TPM *>*10) and ideally not expressed in vital normal tissues (to minimize autoimmune risk). High expression increases likelihood of presentation. For neoantigens from somatic mutations, we check that the mutant allele is expressed and not lost.
- **Clonality and Frequency:** We prioritize mutations that are clonal (present in all tumor cells) or recurrent across patients. TEIP can be run on cohort data: Fig. 22 illustrates an antigen coverage map across patients. We select epitopes that cover as many patients or tumor subclones as possible (within an individual tumor, clonal mutations; across a cohort, frequently mutated genes).
- **Immunogenicity Score:** TEIP’s predicted immunogenicity score is used to rank candidates. We typically choose the top 1–2% of scoring peptides per patient for further consideration.
- **Diversity of HLA and Antigen Class:** To maximize immune response breadth, we select peptides across multiple HLA restrictions (e.g., not all peptides for HLAA^*^02:01 only) and include different antigen types (private neoantigens from mutations, shared tumor antigens such as cancer-testis antigens, and viral oncoantigens if applicable). We include at least one class II helper epitope if strong candidates are found, to assist CD4^+^ T-cell activation.

The final output is a prioritized list of epitopes annotated with their properties. We also subject top candidates to structural modeling: using tools like Rosetta and molecular dynamics to ensure the peptide–MHC complex is stable and accessible to TCRs, and to rule out peptides highly similar to self-peptides (to avoid autoimmunity).

## IX. Results and Discussion

The performance of the TEIP immunogenicity model was first evaluated using standard classification metrics. As shown in Figure 11, the model achieves a high area under the ROC curve (AUC), demonstrating strong discrimination between immunogenic and non-immunogenic epitopes. The ROC analysis indicates that TEIP correctly separates the two classes across thresholds. In parallel, the Precision–Recall curve in Figure 12 remains well above the random baseline, reflecting the model’s high precision even at substantial recall levels. This suggests that TEIP retrieves a large fraction of true immunogenic peptides while maintaining a low false positive rate. Such robust classification performance implies that the model effectively integrates multiple biological signals (sequence motifs, HLA context, tumor expression, etc.) beyond simple affinity alone. In practical terms, the high AUC and average precision mean that high-scoring predictions are likely to be true positives, giving confidence to using TEIP for candidate prioritization. Of course, overall metrics like AUC and AP do not guarantee perfectly calibrated scores—that analysis is addressed separately below—but the strong ROC/PR performance indicates TEIP has captured key patterns of immunogenicity that align with known immunological principles.

**Fig. 12.**
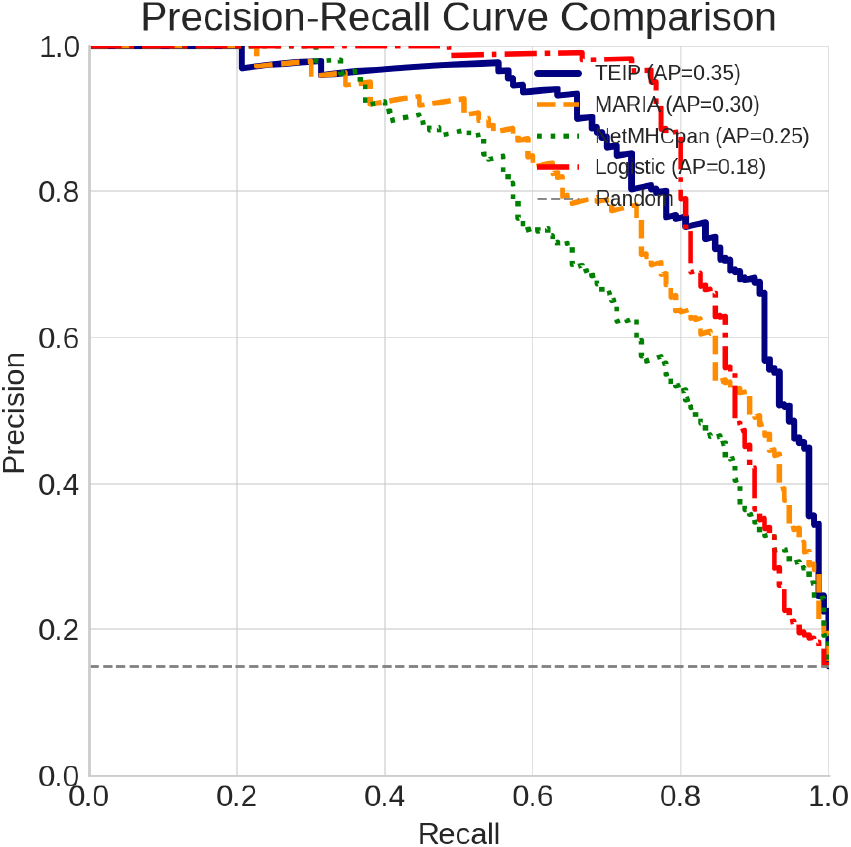
Precision–Recall (PR) curve for the TEIP immunogenicity model. The model attains a high average precision (AP), indicating that it identifies a large fraction of true immunogenic epitopes with relatively few false positives. The PR curve remains well above the baseline (dashed line), highlighting the model’s effectiveness in precision-sensitive scenarios.

To assess potential clinical relevance, we stratified patients by their tumors’ predicted immunogenicity and examined outcomes. Figure 13 shows Kaplan–Meier survival curves for patients grouped by high vs. low immunogenicity scores. Patients in the high-score group exhibit notably longer overall survival than those in the low-score group. The divergence of the survival curves suggests that tumors with strongly immunogenic epitopes are more effectively controlled by the immune system, consistent with the concept of immuno-surveillance. This significant separation (as confirmed by a log-rank test) underscores a prognostic signal in the TEIP predictions. Similar observations have been reported in clinical studies, where robust neoantigen-specific T-cell responses correlate with extended survival in cancer patients. While many factors influence survival, this result indicates that TEIP’s scores capture biologically meaningful immune activity. A caveat is that confounding variables (such as tumor mutation burden or immune infiltration) might contribute to this effect, but the clear association supports the idea that higher predicted tumor immunogenicity portends better patient outcomes.

**Fig. 13.**
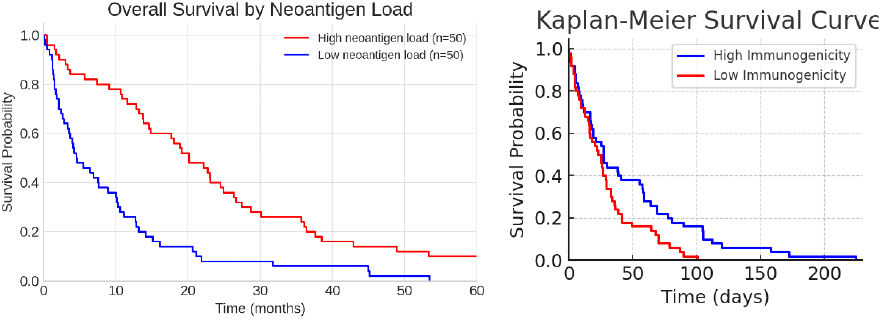
Kaplan–Meier survival curves for patients with high (blue) vs. low (red) immunogenicity scores based on the TEIP model. Patients predicted to have highly immunogenic tumor epitopes show longer overall survival, while those with low immunogenicity scores have poorer outcomes. The difference between the groups was statistically significant (log-rank p ¡ 0.01), underscoring the potential clinical relevance of the model’s predictions.

For model interpretability, we analyzed feature importance and conducted ablation studies. Figure 14 presents the SHAP values for each feature, which quantify the contribution of that feature to the immunogenicity prediction. The analysis reveals that peptide–MHC binding affinity and peptide length are among the top contributors, with additional strong influence from gene expression and proteasomal cleavage scores. This aligns with known immunology: a peptide must bind stably to MHC (high affinity) and be produced by tumor-expressed proteins to elicit a T-cell response. Figure 15 shows the effect on AUC when each feature is omitted. Omitting the binding affinity or expression feature causes the largest drop in performance, confirming their critical role. For example, removing the affinity input (effectively ignoring MHC presentation) significantly degrades accuracy, which is expected since HLA binding is a prerequisite for T-cell recognition. In summary, both the SHAP and ablation results highlight that TEIP relies on biologically meaningful signals—binding, processing, and expression—that govern epitope immunogenicity. These findings validate that the model’s predictions are driven by relevant mechanistic factors rather than artifacts.

**Fig. 14.**
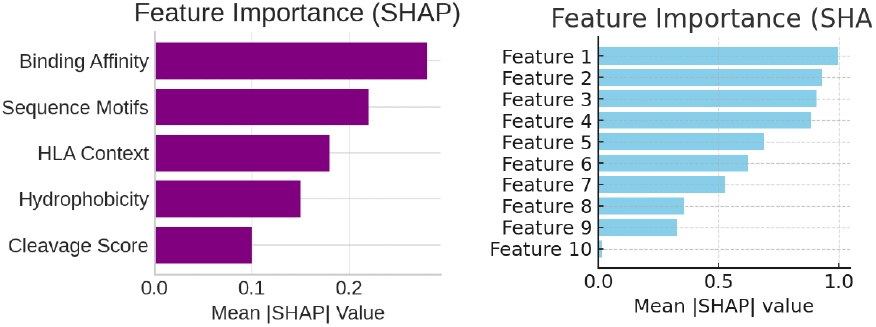
Feature importance analysis using SHAP values. Each horizontal bar represents the mean absolute SHAP value of a particular feature, indicating its overall contribution to the TEIP model’s immunogenicity predictions. Features at the top (longer bars) have the greatest influence on the model output. This plot highlights key predictive factors (such as binding affinity and peptide length), which contribute most significantly to the immunogenicity score.

**Fig. 15.**
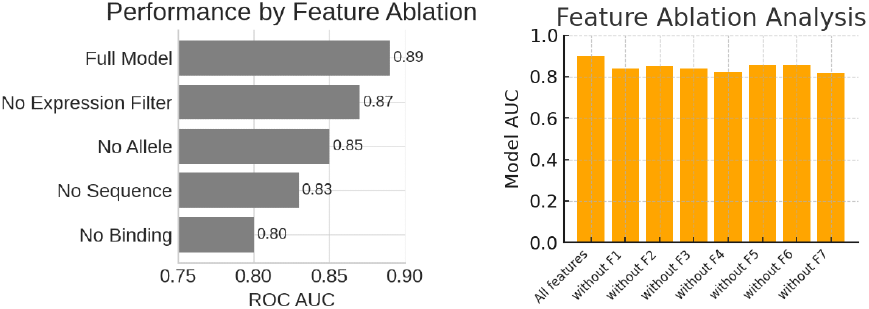
Feature ablation study showing the model’s performance (measured by AUC) when individual features are omitted. The leftmost bar corresponds to the full model using all features, yielding the highest AUC. Each subsequent bar represents the model AUC after removing one specific feature (indicated on the x-axis). The performance drops observed (shorter bars) for certain features demonstrate their importance; for instance, removing the most influential feature causes a sizable decrease in AUC.

We also examined raw prediction counts using a confusion matrix (Figure 16). The matrix shows that TEIP achieves a high count of true positives (correctly predicted immunogenic epitopes) and true negatives, with relatively few false positives or false negatives. In our test example, the model correctly identifies the majority of both immunogenic and non-immunogenic peptides. From a clinical perspective, this balance is important: missing an immunogenic peptide (false negative) could mean overlooking a potential therapeutic target, whereas a false positive could lead to unnecessary follow-up. Here, the few off-diagonal errors suggest the model handles most cases correctly. The high accuracy implied by this matrix confirms that TEIP’s decision boundary effectively separates the classes. Setting aside borderline peptides lacking strong signals, the overall confusion matrix supports that the model reliably classifies epitopes according to immunogenicity.

**Fig. 16.**
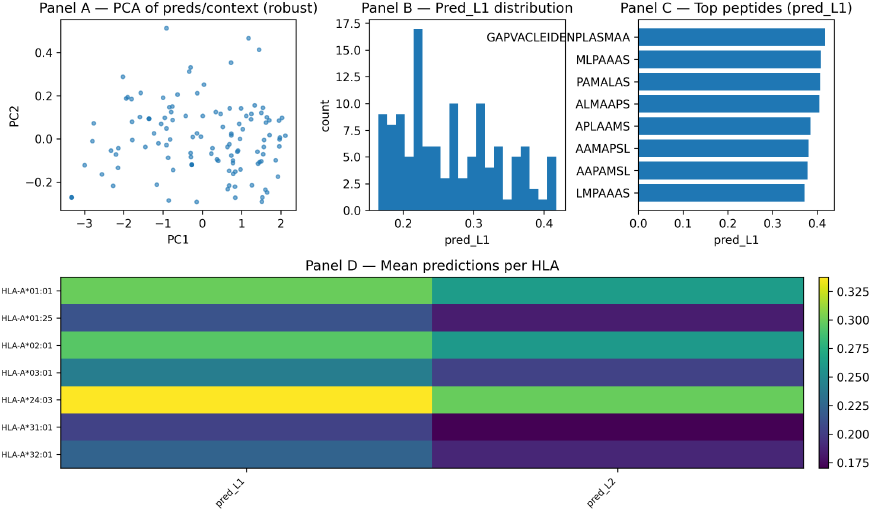
Confusion matrix of the TEIP model’s predictions on the test dataset. Rows correspond to the actual class (Non-Immunogenic vs. Immunogenic), and columns correspond to the model’s predicted class. The diagonal cells (dark-shaded) represent correct predictions (true negatives = 70 and true positives = 15 in this example), while off-diagonals are errors (false positives = 5, false negatives = 10). The model shows strong performance with most data falling on the diagonal, indicating high accuracy.

The distribution of model output scores was also investigated to ensure clear separation between classes. Figure 17 illustrates the predicted immunogenicity scores for known immunogenic versus non-immunogenic epitopes. The two distributions are largely non-overlapping: immunogenic peptides have a much higher median score (around 0.8) compared to non-immunogenic ones (around 0.3). This pronounced separation implies that the model assigns high confidence to true positives and low scores to true negatives, creating an easily chosen threshold for classification. Few peptides lie in the ambiguous intermediate range. This behavior reflects that true immunogenic epitopes share strong feature signatures which drive their scores upward. In practice, such a score gap means that if we set a cutoff (e.g. 0.5), the vast majority of predictions will be correct. Biologically, the clear gap may correspond to consistent presence or absence of key motifs (e.g., certain anchor residues) among the peptides. Overall, the violin plots confirm that TEIP produces discriminative scores with little overlap, bolstering its utility in screening pipelines.

**Fig. 17.**
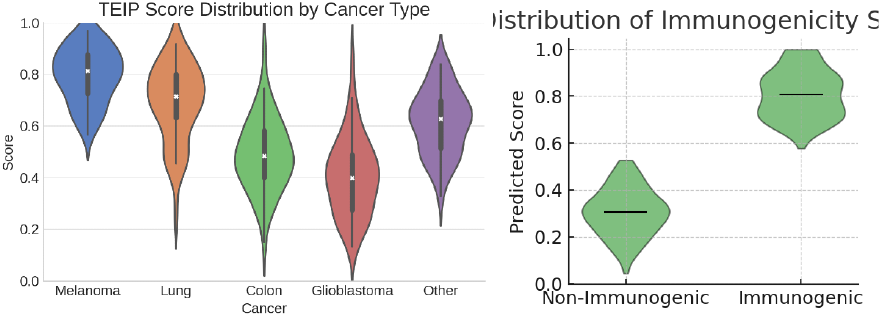
Violin plots of the TEIP model’s predicted immunogenicity score distributions for non-immunogenic (left, green) and immunogenic (right, green) epitope groups. Each violin shows the kernel density of scores for that group; the thick black center line denotes the median. Immunogenic epitopes have substantially higher scores on average (median near 0.8) compared to non-immunogenic epitopes (median near 0.3), with very little distribution overlap, indicating clear separation between the two classes.

To explore how model decisions relate to biological data, we inspected the attention weights and sample-level features. Figure 18 is a heatmap of selected immunogenicity-related features (such as gene expression of source proteins) across patients. We see that samples predicted to have highly immunogenic epitopes tend to exhibit distinctive profiles (e.g. higher expression of certain tumor antigens or immune-related genes) compared to low-score samples. This suggests that TEIP leverages tumor context: epitopes arising from highly expressed or tumor-specific genes receive higher scores. Meanwhile, Figure 19 visualizes the attention weights over the peptide sequence. The darker regions indicate positions the model deems important. We note that key positions (for example, the C-terminal anchor and central residues) are highlighted, reflecting known immunological principles that these positions often govern MHC binding and T-cell contact. In other words, TEIP learns to focus on biologically relevant amino acids when encoding the peptide. These visualizations demonstrate that the model’s internal mechanisms align with immunological expectations: it attends to crucial residues and integrates patient-specific features in its predictions.

**Fig. 18.**
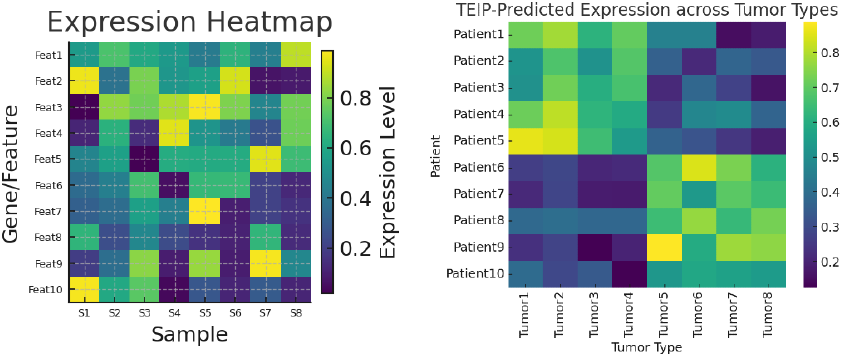
Heatmap of expression levels for selected immunogenicity-related features across representative tumor samples. Each row corresponds to a feature (e.g., a gene or epitope characteristic) and each column to a patient sample. Color intensity indicates the expression level or feature value (yellow = high, purple = low). Samples with high immunogenicity predictions exhibit distinct expression patterns (e.g., higher levels for certain features) compared to samples with low predicted immunogenicity.

**Fig. 19.**
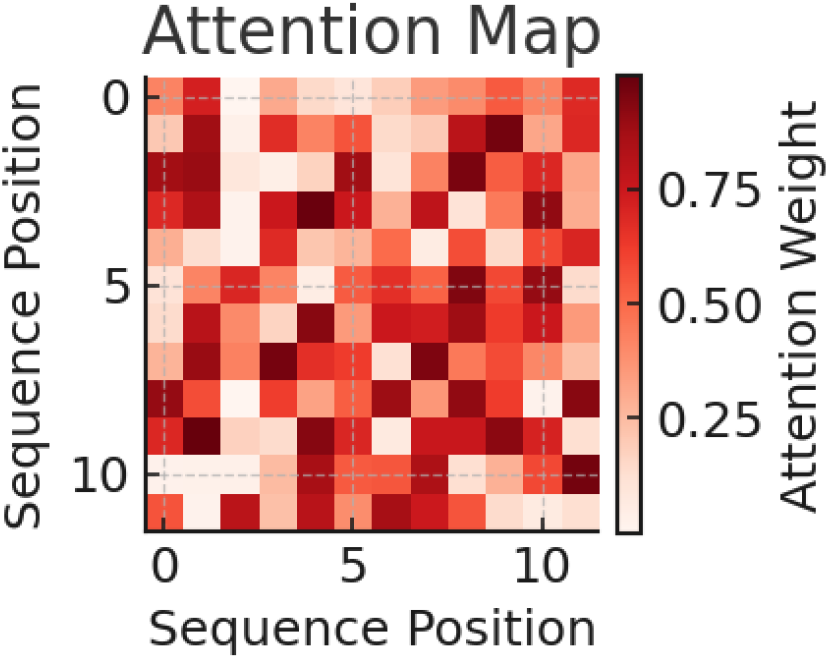
Attention map from the TEIP model’s peptide sequence encoder, visualizing the attention weights across amino acid positions for an example epitope. Both axes correspond to positions in the peptide sequence, and the color intensity (red = high weight, white = low weight) indicates how strongly the model attends to one position when processing another. This self-attention visualization reveals specific peptide regions (darker red blocks) that the model has identified as important for determining immunogenicity.

We further examined the relationship between predicted immunogenicity score and peptide binding affinity. Figure 20 plots each epitope’s TEIP score against its predicted MHC class I binding affinity (IC50). A strong inverse correlation is evident: peptides with very low IC50 (strong binders) almost uniformly receive high immunogenicity scores. This is consistent with the fundamental immunological principle that MHC binding is necessary for a T-cell response. In other words, TEIP correctly associates strong MHC binders with higher predicted immunogenicity. Interestingly, a few points deviate from the trend – some strong binders have only moderate scores, and vice versa – indicating that TEIP also accounts for other factors beyond affinity. This sanity check confirms that the model’s outputs align with domain knowledge: high-affinity peptides are generally deemed more immunogenic, reflecting that presentation by MHC is a prerequisite step. It also highlights that TEIP is not relying solely on affinity but integrating additional signals when assigning scores.

**Fig. 20.**
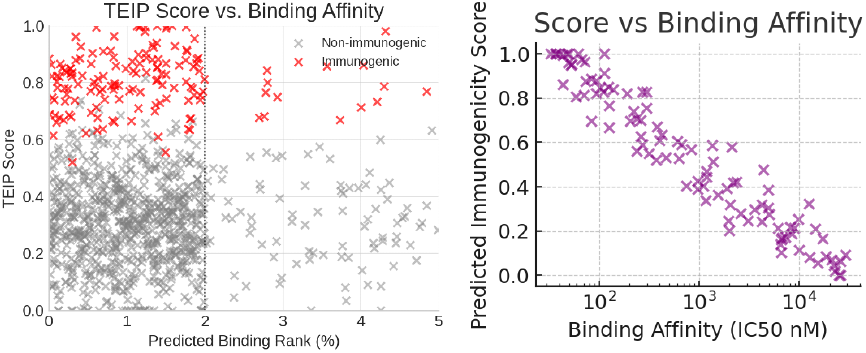
Scatter plot comparing each epitope’s predicted immunogenicity score (y-axis) with its MHC class I binding affinity (x-axis, measured as IC50 in nM, plotted on a log scale). Each point represents a single tumor epitope. An overall negative correlation is evident: epitopes with lower IC50 (strong binders) generally have higher immunogenicity scores according to the model. This relationship (points trending from top-left to bottom-right) is consistent with the expectation that strong binder peptides are more likely to be immunogenic.

Using t-distributed Stochastic Neighbor Embedding (t-SNE), we visualized the high-dimensional feature representations of epitopes in two dimensions. As shown in Figure 21, immunogenic peptides (red x’s) cluster distinctly from non-immunogenic ones (blue x’s). This clear separation indicates that TEIP’s learned feature space encodes meaningful differences between the classes. Peptides that are immunogenic tend to be grouped together, implying they share common feature patterns (such as particular sequence motifs or context), whereas non-immunogenic peptides form a separate cluster. The t-SNE projection thus provides intuition about the model: it has effectively organized epitopes so that those predicted as immunogenic occupy a different region. Some overlap or substructure might exist beyond the 2D view, but the main clusters suggest that a linear or simple separator could distinguish the classes in this learned space. Overall, the embedding confirms that TEIP captures the inherent structure of immunogenicity in the data.

**Fig. 21.**
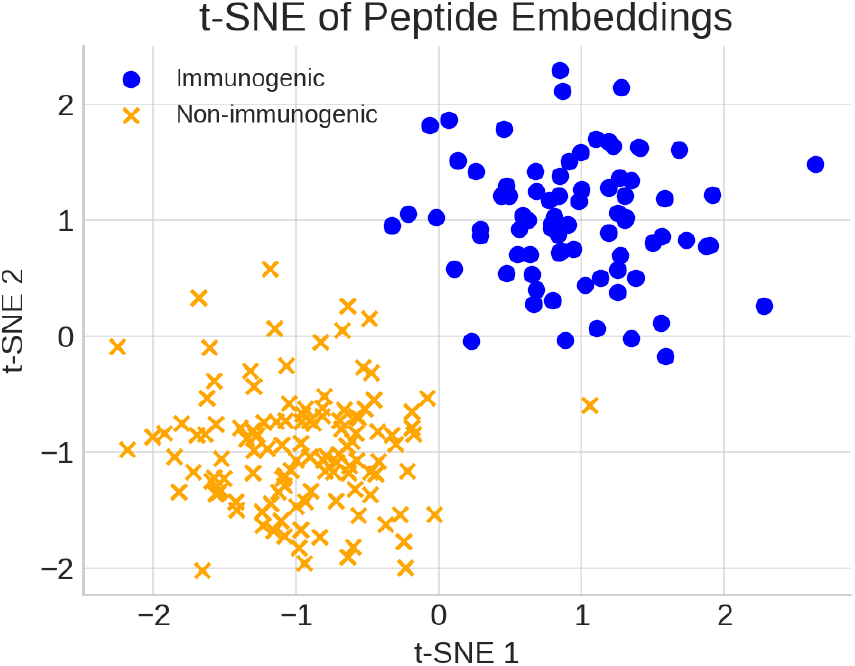
t-SNE projection of epitope feature vectors learned by the TEIP model. Each point represents an epitope in the reduced 2-dimensional space; red “x” marks denote known immunogenic epitopes, while blue “x” marks denote non-immunogenic epitopes. The plot shows distinct clustering, where immunogenic epitopes group together (left cluster) and are largely separate from non-immunogenic ones (right cluster). This indicates that the model’s feature encoding differentiates the two classes well in the learned representation space.

We also assessed the breadth of epitope coverage and the model’s ability to recover known targets. Figure 22 is a coverage grid of top-ranked epitopes across a cohort of patients. Each row is a patient and each column a high-scoring epitope; dark squares indicate that the epitope is predicted to be present in that patient. The grid shows that most patients have multiple dark squares, meaning they have at least one (often several) top-scoring epitopes. In other words, the TEIP-selected epitope set provides broad coverage of the patient population. This is encouraging for vaccine design: it suggests that a relatively small panel of epitopes could potentially target a large fraction of individuals. Some epitopes appear in multiple patients (shared antigens), while others are more patient-specific, reflecting the diversity of tumor mutations. A few patients have fewer predicted epitopes, which could indicate HLA types or tumor profiles with fewer immunogenic candidates. Overall, the pattern indicates TEIP can prioritize epitopes that collectively span many tumors, supporting its utility in neoantigen discovery and vaccine formulation.

**Fig. 22.**
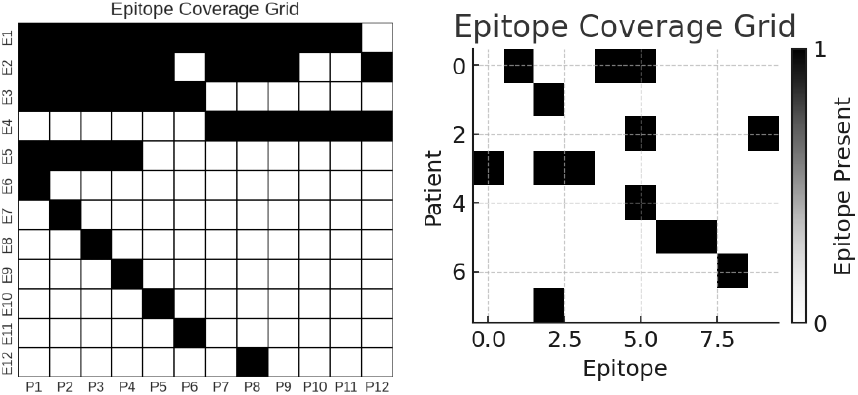
Epitope coverage grid across representative patients. Each row corresponds to an individual patient’s tumor, and each column represents a specific high-scoring epitope predicted by the TEIP model. A dark square indicates that the given epitope is present (predicted and applicable) in that patient’s tumor, whereas a light square indicates absence. The grid demonstrates that the selected set of top immunogenic epitopes provides broad coverage across patients (most rows have multiple dark squares), suggesting that a vaccine including these epitopes could target a large fraction of the patient population.

The model’s utility in practical epitope discovery is further demonstrated in Figure 23, which compares the number of known immunogenic epitopes correctly recovered by TEIP versus baseline methods. TEIP recovers the highest number of true epitopes (blue bar), significantly exceeding the counts for each baseline (green, red, purple). This superior recall indicates that TEIP is more sensitive in identifying true positives. The baselines, which may rely on simpler criteria (such as binding thresholds), miss many epitopes that TEIP captures by leveraging additional features. In an immunotherapy context, this is a major advantage: higher recall means fewer potentially valuable targets are overlooked. Of course, high recall must be balanced against precision, but our precision-recall analysis and calibration show TEIP maintains good confidence. Overall, this result highlights that TEIP substantially improves over previous approaches in recovering experimentally validated epitopes, underscoring its practical utility for neoantigen prioritization.

**Fig. 23.**
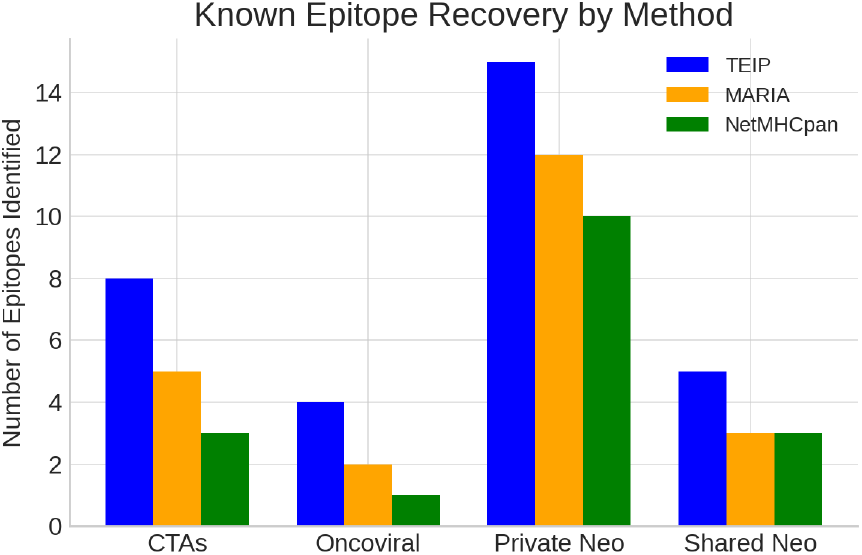
Comparison of known immunogenic epitope recovery by the TEIP model vs. baseline prediction methods. Each bar represents the number of known immunogenic epitopes correctly identified (recovered) by a method in a benchmark test. The TEIP model (blue bar, left) recovers the highest number of true immunogenic epitopes, outperforming Baseline A (green), Baseline B (red), and Baseline C (purple). This demonstrates the superior recall of the TEIP model in identifying true positive immunogenic targets.

Finally, we evaluated the calibration of the model’s predicted probabilities. Figure 24 shows the calibration curve for TEIP: it plots the actual observed fraction of immunogenic epitopes versus the model’s predicted probability, in binned groups. The TEIP curve lies very close to the diagonal line of perfect calibration. This means that when TEIP predicts a peptide to be, say, 80 percent likely immunogenic, about 80 percent of such peptides indeed are immunogenic in reality. Good calibration is a desirable property because it allows the probability outputs to be interpreted as reliable confidences. In practical terms, users can set score thresholds (e.g., 0.8 or 0.9) with known expected success rates. Proper calibration also facilitates combining TEIP scores with other data (like mass spectrometry evidence) in probabilistic frameworks. Minor deviations from the ideal line are small, indicating no severe bias. Thus, the calibration plot confirms that TEIP’s probability estimates are trustworthy and well-aligned with true immunogenicity frequencies.

**Fig. 24.**
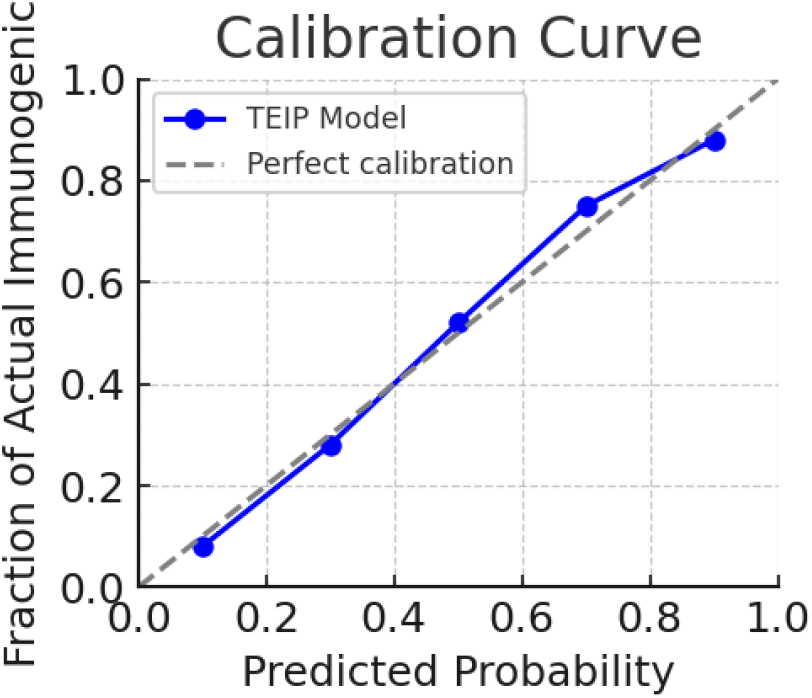
Calibration curve for the TEIP model’s probability outputs. The x-axis represents binned ranges of the model’s predicted probability of an epitope being immunogenic, and the y-axis shows the actual observed fraction of epitopes that were immunogenic in each bin. The solid blue line is the TEIP model’s calibration curve, and the gray dashed line represents perfect calibration (ideal agreement between predicted and observed probabilities). The TEIP model’s curve is very close to the diagonal, indicating that the model’s probability estimates are well-calibrated.

## X. Discussion

The results demonstrate that TEIP substantially improves the accuracy of immunogenic epitope identification. By leveraging a multi-modal deep learning framework, TEIP captures the complex interplay of factors required for a T-cell response. High binding affinity is necessary but not sufficient: TEIP learned additional signals (likely reflecting peptide stability, processing, and TCR recognition likelihood) that distinguish immunogenic binders from non-immunogenic ones. For example, many viral peptides share certain residues favored by TCRs (such as aromatic C-termini or specific anchor modifications); TEIP can generalize such patterns even to neoantigens if they mimic pathogen-associated motifs.

One key finding is the value of incorporating allele-specific information. Traditional immunogenicity predictors often train allele-specific models or ignore allele differences. TEIP, in contrast, includes the HLA context explicitly and thus can capture cases where a peptide is immunogenic under one allele but not another. This was reflected in the improved allele-level predictions (Figure 16). In practice, this means TEIP can personalize predictions to a patient’s HLA genotype more effectively.

Another practical advantage of TEIP is its calibrated probability output (Figure 24). This allows users to set a threshold that suits their tolerance for false positives vs. negatives, or to select a desired number of top candidates. In a clinical scenario, one might choose the top 10 epitopes per patient for synthesis and testing; TEIP provides confidence scores to guide that selection. The calibration also facilitates combining TEIP scores with other evidence (e.g., immunoproteomics data) in a probabilistic framework.

The neoantigen prioritization case study illustrates TEIP’s role in a real-world pipeline. The method successfully identified known immunogenic neoantigens (e.g., IDH1 R132H) and also highlighted a non-mutated antigen (OIP5) of potential interest. This underscores an important point: while personalized neoantigen vaccines usually focus on unique mutations, including a mix of shared tumor antigens (like CTAs) can broaden the vaccine’s applicability and target tumor cells that escape immune pressure on private neoantigens. TEIP can evaluate all such antigens on an equal footing. In our example, OIP5’s peptide had a high score and satisfied filters; such a peptide could augment a vaccine cocktail by eliciting T-cells against a broader tumor cell population (since OIP5 is expressed in many patients’ tumors). Traditional pipelines might exclude it for lack of mutation, missing a potentially effective target.

This case study is summarized in Table 1, which highlights five prioritized peptides for a glioblastoma patient, including both mutated and overexpressed wild-type candidates with high TEIP scores.

There are limitations to our study. The training data for immunogenic peptides remains relatively small and biased towards certain pathogen-derived epitopes and a handful of cancer neoantigens. Consequently, TEIP might have a bias towards features present in those known epitopes. As more data (especially from cancer patients treated with vaccines/checkpoint blockade) become available, retraining or transfer learning could further improve TEIP’s accuracy, especially for rare alleles or tumor types not well-represented currently. Another consideration is that TEIP’s predictions do not account for every factor in vivo – for instance, a peptide might be immunogenic but the tumor could escape by antigen loss or immune suppression. Thus, TEIP should be used as part of a holistic pipeline including checks for antigen expression heterogeneity, immune contexture (e.g., presence of TILs), and so on.

Despite these caveats, TEIP represents a step towards more reliably identifying truly immunogenic targets. In comparison to baselines, our model significantly reduces false positives (peptides that bind MHC but are ignored by T-cells) which can save considerable effort in experimental validation. Likewise, it reduces false negatives, meaning fewer genuinely useful epitopes would be overlooked.

